# The frailty syndrome as an emergent state of parallel dysregulation in multiple physiological systems

**DOI:** 10.1101/2020.07.22.214718

**Authors:** Ahmed Ghachem, Linda P. Fried, Véronique Legault, Karen Bandeen-Roche, Nancy Presse, Alan A. Cohen

**Affiliations:** Research Center on Aging, CIUSSS-de-l’Estrie-CHUS, Sherbrooke, QC, Canada; Mailman School of Public Health and Columbia University Medical Center, New York, NY, United States; Groupe de recherche PRIMUS, Department of Family Medicine, University of Sherbrooke, Sherbrooke, QC, Canada; Department of Biostatistics and Center on Aging and Health, Bloomberg School of Public Health, The Johns Hopkins University, Baltimore, MD, United States; Centre de Recherche de l’Institut Universitaire de Gériatrie de Montréal (CRIUGM), Montréal, QC, Canada; Faculty of Medicine and Health Sciences, Université de Sherbrooke, Sherbrooke, QC, Canada

**Keywords:** Aging, Complexity, Emergence, Homeostasis, Phenotypic frailty, Physiological Dysregulation, Resilience

## Abstract

Frailty is a clinical syndrome often present in older adults and characterized by a heightened vulnerability to stressors. The biological antecedents and etiology of frailty are unclear despite decades of research: frailty is associated with dysregulation in a wide range of physiological systems, but no specific cause has been identified. Here, we test predictions stemming from the hypothesis that there is no specific cause: that frailty is an emergent property arising from the complex systems dynamics of the broad loss of organismal homeostasis. Specifically, we use dysregulation of six physiological systems using the Mahalanobis distance approach in two cohorts of older adults to test the breadth, diffuseness, and nonlinearity of associations between frailty and system-specific dysregulation. We find clear support for the breadth of associations between frailty and physiological dysregulation: positive associations of all systems with frailty in at least some analyses. We find partial support for diffuseness: the number of systems or total amount of dysregulation is more important than the identity of the systems dysregulated, but results only partially replicate across cohorts. We find partial support for nonlinearity: trends are exponential but not always significantly so, and power is limited for groups with very high levels of dysregulation. Overall, results are consistent with – but not definitive proof of – frailty as an emergent property of complex systems dynamics. Substantial work remains to understand how frailty relates to underlying physiological dynamics across systems.

## Introduction

Frailty is a distinct medical geriatric syndrome defined by the presence of three or more of the following criteria: unintentional weight loss, low physical activity, slowed motor performance, low energy, or low strength (Fried et al. 2001a). Studies have shown that frailty is associated with multiple adverse health problems and significantly increases risk of falls, institutionalization, disability, and mortality (Fried et al. 2001a; Clegg et al. 2013). The etiology of frailty is still, however, not clear. Previous studies have suggested that dysregulation of biomarkers could be involved in the development of frailty, and indeed, cohort studies have shown that levels of pro-inflammatory cytokines including interleukin-6, C-reactive protein and tumor necrosis factor-α are higher among frail older adults (Leng et al. 2002; Qu et al. 2009). High levels of other biomarkers such as neopterin, a biomarker of immune system activation, total white blood cell count, growth hormone, IGF-1, cortisol and vitamin D were also significantly associated with frailty risk in community-dwelling older adults (Leng et al. 2007, 2009; Hubbard et al. 2008; Shardell et al. 2009). Taken together, these findings suggest a close relationship between underlying aging-related biological processes and the frailty syndrome. However, it is not yet clear whether biomarker changes cause frailty or whether frailty tends to co-occur with other aging-related conditions to cause changes in biomarkers levels.

Frailty has been conceived of as an emergent property of a broad underlying physiological state, rather than as a dysregulation of a single biomarker or even of a series of independent biomarkers (Fried et al. 2009, 2020). Under this view, frailty is not the product of a specific physiological or biochemical pathway, but rather emerges from the complex systems dynamics that maintain an organism in homeostasis. Loss of this homeostasis (dysregulation) is thought to occur progressively with age (Seplaki et al. 2005; Li et al. 2015; Cohen 2016), and frailty could be the emergent manifestation of the end-stage loss of homeostatic resilience and the accompanying decline in ability to respond successfully to stressors (Nakazato et al.; Olde Rikkert et al. 2016; Gijzel et al. 2017). This conception of frailty makes a number of predictions, few of which have been rigorously tested and replicated. Notably but not exhaustively, frailty should (a) reflect a broad loss of resilience (not just loss of resilience to one or several types of stressors); (b) emerge from compromised network dynamics implying dysregulation of multiple physiological systems; (c) present a series of coherent clinical signs and symptoms even when the underlying physiological mechanisms are diverse, differing from one individual to another; and (d) have a non-linear relationship with its physiological antecedents, emerging rather abruptly as a certain threshold of underlying dysregulation is achieved. Here, we wish to test these last three propositions, the *breadth, diffuseness* and *non-linearity* of the physiological antecedents of frailty.

The hypothesis that frailty is the result of an age-related multi-system physiological dysregulation (PD) has been previously evoked, and was specifically theorized and investigated by Fried and al. (2009). Using one or two biomarkers per system and *a priori* clinical knowledge to calculate dysregulation, results of this study showed, first, a non-linear and positive relationship between the number of abnormal physiological systems including anemia, endocrine function, inflammation, micronutrients, adiposity, fine motor status and risk of frailty. Second, older women with three or more abnormal systems were more likely to be frail, with increasing odds of frailty with increasing number of abnormal systems, even after controlling for individual system, number of chronic diseases, and chronological age (Fried et al. 2009).

Recently, we developed and validated, in different species and human populations, a novel and rigorous approach to measure PD based on the Mahalanobis statistical distance (Cohen et al. 2013, 2014, 2015a; Milot et al. 2014; Li et al. 2015; Dansereau et al. 2019). Our previous results showed that dysregulation of 37 biomarkers, grouped into six standard physiological systems, significantly increases with age and predicts multiple health outcomes including cardiovascular diseases, frailty, and mortality (Li et al. 2015). Crucially, the dysregulation levels of these systems are only very weakly correlated with each other after controlling for age, implying that dysregulation proceeds nearly independently in each system.

Here, we attempt to replicate the findings of Fried et al. (2009), with two key differences. First, we are using a different list of physiological systems, that derived based on the Mahalanobis distance approach, and thus agnostic to physiological theory about what systems might drive frailty. This difference is crucial because it allows us to assess whether the specific choice of systems is necessary, or whether frailty might be seen as emerging from a more general physiological “soup.” Second, in addition to using the Women’s Health and Aging Study (WHAS), we also use the Quebec Longitudinal Study on Nutrition and Successful Aging (NuAge) study to provide cross-population validation. We tested three main hypotheses. First, we hypothesized that parallel dysregulation of multiple physiological systems drives the development of frailty (breadth of frailty’s physiological antecedents). We predicted that frailty risk would increase with higher physiological dysregulation in all or most systems, and this would be true of the individual frailty criteria as well (minimal specificity between dysregulation systems and frailty criteria). Second, we hypothesized that frailty would not depend on which systems were dysregulated as much as how many systems were dysregulated (diffuseness). Third, we hypothesized that frailty would be non-linearly associated with dysregulation levels (non-linearity). If frailty is an emergent property of a dysregulated complex system, frailty risk should not increase linearly with PD level; rather, there should be an exponential relationship or a threshold effect. The relationship between frailty risk and PD should be nonlinear, with a sharp increase of frailty risk at a given number of dysregulated systems. For this hypothesis, we made predictions on two levels. First, within each system, discriminatory power to diagnose/predict frailty should increase with higher levels of dysregulation. Second, frailty risk should increase non-linearly as a function of the number of systems dysregulated. Together these predictions will help test the broader hypothesis that systems become increasingly dysregulated with aging, and that it is the critical mass of dysregulated systems that drives the development of frailty.

## Methods

### Study populations

We used data from WHAS I and II, and from the NuAge study. WHAS sampling and data collection procedures have been described in detail elsewhere (Simonsick et al. 1997; Fried et al. 2001b). Briefly, WHAS I and II are two studies which investigated health and physical functioning in community-dwelling older women aged 65 years and older. Women from WHAS I represent the one-third most disabled older women living in the community, and selection criteria included a Mini-Mental State Examination (MMSE) score of greater than 18 and difficulty in two or more of the following functional domains: mobility, upper extremity, household management tasks, and self-care tasks. On the other hand, women from WHAS II had either no difficulty or difficulty in only one functional domain, and MMSE scores of 24 or greater. We performed analyses on WHAS I and WHAS I and II combined, including inverse probability weights using the “survey” package in R, and accordingly limited observations to where age distributions overlapped. We present some stratified analyses in the Supplement, but encourage caution in their interpretation, as the sampling for each separate cohort was based on health state and thus strongly associated with frailty.

The NuAge cohort consists of 1,793 mostly Caucasian community-dwelling men and women assessed as being in good general health at recruitment and having between 68-86 years (Gaudreau et al. 2007). Participants were identified via a sex-and age-stratified random sample from a population-wide health insurance list, and were included if they were free of disabilities in activities of daily living, had no cognitive impairment (Modified MMSE score > 79), and were able to walk one block or to climb one flight of stairs without rest. Participants were re-examined annually for 3 years. A total of 1,754 (97.8%) participants agreed for the integration of their data and biological samples into a research bank, now called the NuAge Database and Biobank, whom we considered in our analyses.

### Biomarker selection

Biomarkers were selected based on availability and adequate sample sizes. A total of 33 biomarkers in WHAS and 28 in NuAge, grouped into respectively six and five previously validated physiological systems (Li et al. 2015), were used to calculate PD, globally and per system. Due to differences in availability of biomarkers across datasets, the precise choice of biomarkers included in some systems vary slightly (e.g. oxygen transport and micronutrients). However, our previous work has shown that PD produces a robust signal independent of precise biomarker composition (Cohen et al. 2015a). Sample sizes also varied across systems due to differences in biomarker availability (Table S2). As mentioned above, we used the full list of PD systems previously validated, agnostic to specific theory about which should relate to frailty.

After excluding individuals having missing data in biomarkers and measures of functioning, for NuAge, the sample size was highly variable across systems (Table S2). This difference in sample size is due to a more extensive serum biomarker analysis conducted in 2016 on a subsample of ∼750 individuals that were selected at random among 904 of the 1754 participants who met the following criteria: (1) the individual needed to have blood sampling conducted without any missing intermediate visits; (2) the individual needed at least two visits with blood samples; and (3) there had to be a sufficient number of stored aliquots at all visits to analyze the selected biomarkers.

Details for laboratory assessment of biomarkers in WHAS have been described elsewhere (Mielke et al. 2008; Semba et al. 2010). In NuAge, biomarkers were analyzed in clinical laboratories using standardized and well-established methods (Biron, Inc., Brossard, Qc; Centre Hospitalier de l’Université de Sherbrooke; Centre Hospitalier de l’Université de Montréal). Tocopherols and β-carotene were analyzed following the protocol described in Khalil et al (Khalil et al. 2011).

### Calculation of Physiological dysregulation

PD was calculated using the Mahalanobis distance “DM” based on the following formula:

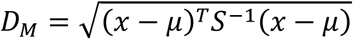

where x is the vector of biomarker values for an individual, μ is the equal-length vector of reference population means for each biomarker and S is the reference population variance-covariance matrix for the biomarkers (Mahalanobis 1936).

PD was calculated by system and globally using all biomarkers for each participant at each time point in the two datasets. Briefly, DM measures how aberrant or unusual an individual’s biomarker profile is compared to a reference population. Here, we used one random visit per individual as the reference population, for each dataset (or subset). Numerous previous studies have shown that a high PD score indicates a worse biomarker profile and higher risk of adverse outcomes in diverse populations (Kraft et al. (in press); Cohen et al. 2014; Arbeev et al. 2016; Belsky et al. 2018). All biomarkers were first log- or square-root-transformed as needed to approach normality (see Table S1 for details) and then were standardized (centered at the mean of the reference population and divided by the standard deviation of the reference population). Presence or absence of dysregulation in each system was dichotomized for some analyses by considering dysregulation to be present if the system score was above the 75^th^ percentile for the respective dataset.

### Frailty phenotype

Fried’s frailty criteria were used to identify the frailty phenotype, adapted to each respective dataset (Table S3). The presence of three or more of these criteria indicates clinical frailty syndrome and one to two criteria indicates a pre-frail state. Those meeting none of the criteria were considered as non-frail (Fried et al. 2001a). When individuals had missing values for some criteria, we used the sum of available criteria to define the frailty phenotype. Correcting for missing criteria, i.e. dividing the sum by the number of available criteria, did not significantly affect the results (data not shown).

### Statistical analysis

Continuous data are presented as mean ± SD. Age was calculated to the nearest day at each visit. PD scores were natural log transformed (except when specified otherwise) and also centered to zero, and standardized to one standard deviation. All analyses were performed using R versions 3.5.0 and 4.0.0, with additional packages “lme4” v1.1-23 (Bates et al. 2015), “survey” v4.0 (Lumley et al. 2020), “ordinal” v2019.12-10 (Christensen 2019), “splines” v4.0.0 (Bates and Venables), and “factoextra” v1.0.7 (Kassambara and Mundt 2020). We do not conduct formal significance tests (i.e., we do not subscribe to the frequentist hypothesis testing framework) (Leek et al. 2017), but we informally refer to results as “significant” when *p* < 0.05 for ease of discussing trends across large numbers of analyses. Both datasets contained multiple datapoints per individual; here, we conducted cross-sectional analyses using all data points, controlling for individual with a random effect in mixed effects models.

### Biomarker profiles according to frailty phenotype

To assess how frail individuals’ biomarker profiles deviate compared to pre-frail and non-frail, Principal Component Analyses (PCA) were performed on the full set of biomarkers for each physiological system using svyprcomp function (survey package) for WHAS and prcomp function for NuAge. The first and second axes that explained the most variance in all the biomarkers were plotted using the PCA-biplot by frailty phenotypes. Results are presented with 95% ellipse to show individuals’ dispersion for each frailty phenotype.

### Association between physiological dysregulation level and frailty risk

First, to assess the association between frailty phenotype and PD score, we performed logistic regression models with PD as a fixed effect and both pre-frail and non-frail phenotypes included in the reference group. We also assessed frailty risk as a function of PD with proportional odds models using all three categories, i.e. non-frail, pre-frail, and frail (clm and clmm functions, ordinal package, or svyolr function, survey package). Relationships between PD score and total number of frailty criteria were assessed using Poisson regression models. Lastly, we used logistic regression to assess relationships between individual frailty criteria and PD scores. All models controlled for age using a cubic spline (bs function, fda/splines packages) and included the individual as a random effect when longitudinal measures were available. Sex was controlled for in analyses on NuAge.

### Associations between frailty and number of dysregulated systems

Using the 75^th^ percentile as mentioned above, we assigned each observation a number of dysregulated systems, excluding global PD (0-6 in WHAS and 0-5 in NuAge). In order to ensure sufficient sample size in each bin for visualizations, we regrouped these into 0 vs 1-2-3 vs 4+ dysregulated systems. We also used an alternative model with a sum of dysregulation in each system excluding global, weighted by the inverse of the number of biomarkers included in each system in order to ensure equal weight for each system. We then assessed associations between number/sum of dysregulated systems and frailty state (non-frail, pre-frail and frail) visually. We also ran logistic and proportional odds models to assess frailty risk, controlling for which systems were dysregulated and for age as a cubic spline. This model reciprocally assessed whether the presence of any individual dysregulated system increased risk of frailty net of the number of systems, and whether the number of systems predicted frailty net of the identity of the systems. Because there are very few frail individuals in the NuAge cohort (3.8% of observations), we combined pre-frail with frail to increase our statistical power, whereas pre-frail was combined with non-frail in WHAS for the logistic regressions.

### Assessment of nonlinearity

We assessed nonlinearity of associations between frailty and PD in several ways. First, visually. Second, by modelling frailty risk with logistic regression models as a quadratic function of dysregulation in each system. Third, we used linear regression with the sum of dysregulation scores across systems (weighted to have equal representation). We modelled this sum as a function of frailty status in three ways. First, with frailty as an unordered categorical variable (non-frail, pre-frail, frail; this model does not test for nonlinearity). Second, with frailty as a numeric variable with three levels, 0, 1, and 2 respectively, and with a quadratic term. Third, with frailty as a numeric variable (0, 1, and 2), and with an additional categorical variable to distinguish frail from pre-frail and non-frail. This last model assesses the additional increase in dysregulation with frailty status, compared to what would be expected if the effect were linear across frailty statuses.

## Results

Demographic characteristics of our populations are shown in Table 1. Note the small numbers of frail individuals in WHAS II and in NuAge, particularly sex-stratified. Based on these results, we decided to present results for NuAge combined but not stratified by sex in the main text, with stratified analyses in the supplement along with the two separate WHAS cohorts. For the same reason, while we prefer to group prefrail with non-frail in dichotomizing, for some subgroup analyses we have put prefrail with frail to increase statistical power.

**Table 1.**
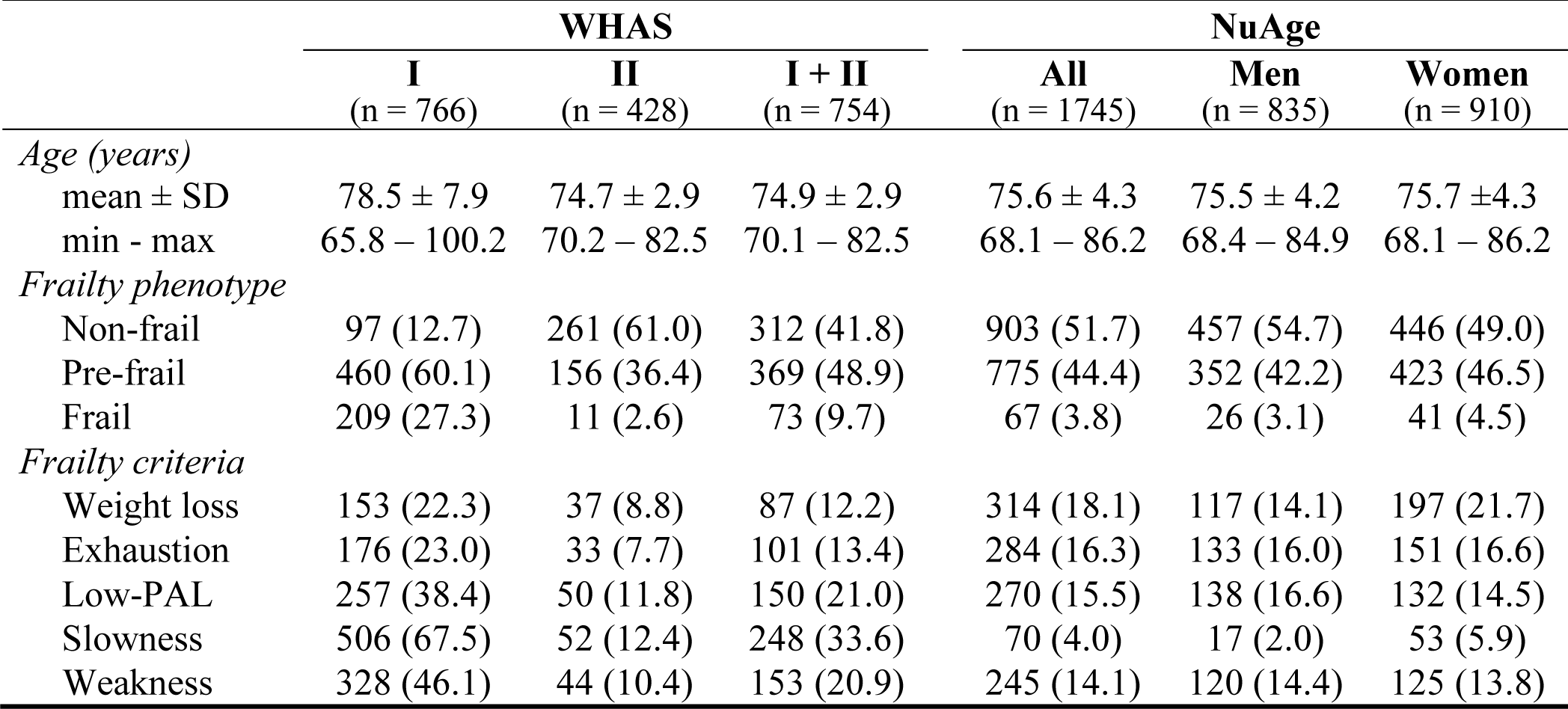
Characteristics of participants at first visit used, indicated as numbers (with percentages in parenthesis), unless specified otherwise.

|  | WHAS |  |  | NuAge |  |  |
| --- | --- | --- | --- | --- | --- | --- |
|  | I<br>(n = 766) | II<br>(n = 428) | I + II<br>(n = 754) | All<br>(n = 1745) | Men<br>(n = 835) | Women<br>(n = 910) |
| *Age (years)* |  |  |  |  |  |  |
| mean ± SD | 78.5 ± 7.9 | 74.7 ± 2.9 | 74.9 ± 2.9 | 75.6 ± 4.3 | 75.5 ± 4.2 | 75.7 ± 4.3 |
| min - max | 65.8 – 100.2 | 70.2 – 82.5 | 70.1 – 82.5 | 68.1 – 86.2 | 68.4 – 84.9 | 68.1 – 86.2 |
| *Frailty phenotype* |  |  |  |  |  |  |
| Non-frail | 97 (12.7) | 261 (61.0) | 312 (41.8) | 903 (51.7) | 457 (54.7) | 446 (49.0) |
| Pre-frail | 460 (60.1) | 156 (36.4) | 369 (48.9) | 775 (44.4) | 352 (42.2) | 423 (46.5) |
| Frail | 209 (27.3) | 11 (2.6) | 73 (9.7) | 67 (3.8) | 26 (3.1) | 41 (4.5) |
| *Frailty criteria* |  |  |  |  |  |  |
| Weight loss | 153 (22.3) | 37 (8.8) | 87 (12.2) | 314 (18.1) | 117 (14.1) | 197 (21.7) |
| Exhaustion | 176 (23.0) | 33 (7.7) | 101 (13.4) | 284 (16.3) | 133 (16.0) | 151 (16.6) |
| Low-PAL | 257 (38.4) | 50 (11.8) | 150 (21.0) | 270 (15.5) | 138 (16.6) | 132 (14.5) |
| Slowness | 506 (67.5) | 52 (12.4) | 248 (33.6) | 70 (4.0) | 17 (2.0) | 53 (5.9) |
| Weakness | 328 (46.1) | 44 (10.4) | 153 (20.9) | 245 (14.1) | 120 (14.4) | 125 (13.8) |

### Biomarker profiles by frailty phenotype

We plotted PCA biplots of biomarkers by system, color-coded by frailty phenotype (Fig. 1 and Figs S1 and S2). Overall, frail individuals had greater dispersion around the centroid, suggesting greater biomarker dysregulation, compared to pre-fail and non-frail. In some cases, this was simply greater dispersion, whereas in others the dispersion was directed toward a certain profile. For example, for oxygen transport biomarkers, frail individuals displayed lower values of hemoglobin, hematocrit and iron, reflecting anemia. For lipid biomarkers, frail individuals had lower HDL-C levels and higher triglyceride levels and cholesterol-HDL-C ratios compared to non-frail. Overall, results were highly consistent across systems, suggesting that frail individuals had different biomarker profiles compared to pre-frail and non-frail. However, there was still marked overlap between the frail, pre-frail, and non-frail individuals, indicating that there is no clear or universal biomarker signature of frailty among the measured biomarkers.

**Figure 1.**
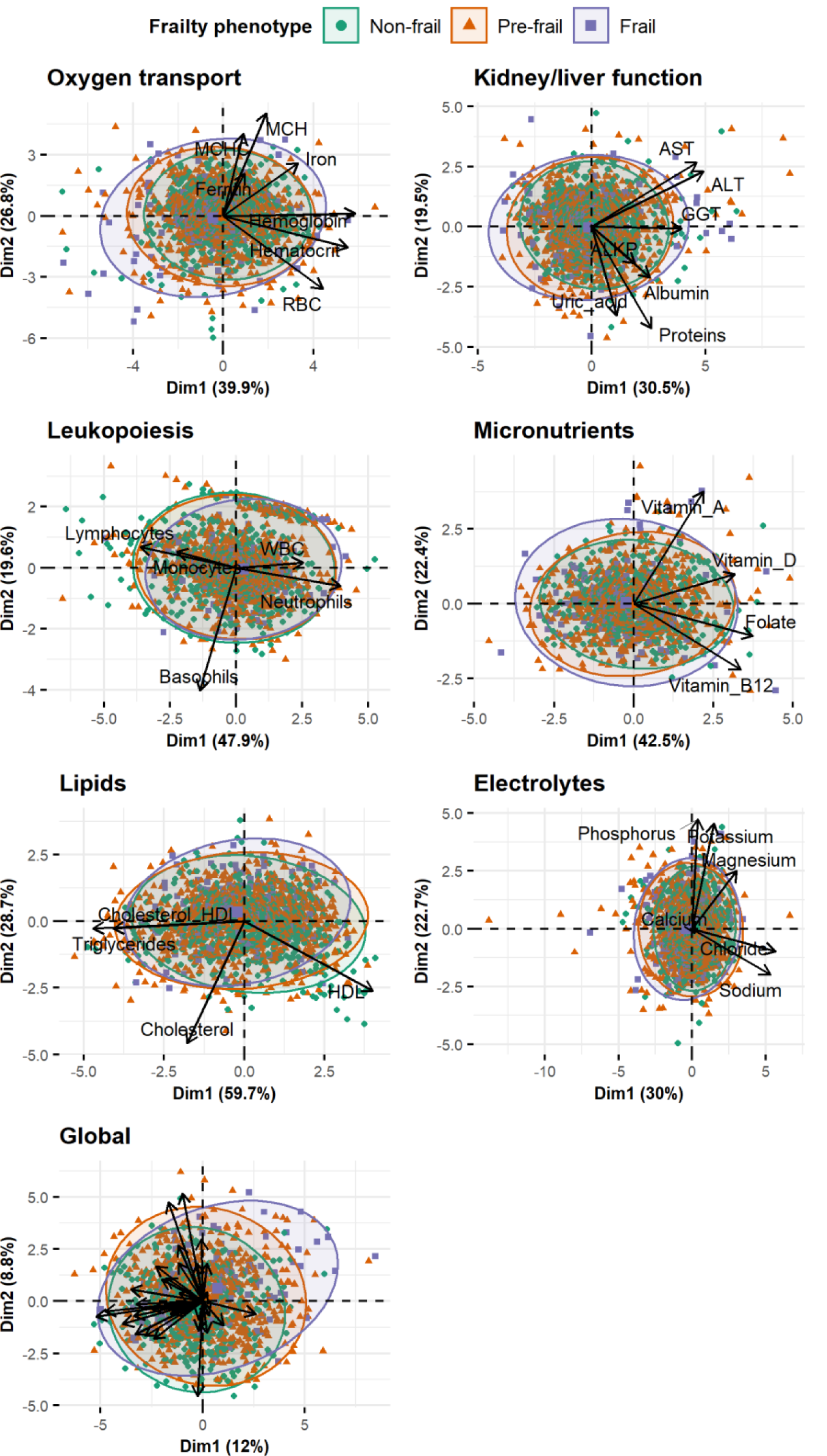
Principal component analyses to characterise biomarkers profile of non-frail *vs* pre-frail *vs* frail individuals in WHAS (I + II) for each physiological system and globally. Individuals (points) together with 95% ellipses (dispersion) by frailty phenotypes for each physiological system are presented. Different colors indicate frailty phenotypes. See Figs. S1-S2 for WHAS I and NuAge results.

### Physiological dysregulation level according to frailty phenotypes

Overall, frail individuals displayed higher PD levels compared to pre-frail and non-frail, globally and for most systems, in both datasets (Fig. 2, Fig. S3). Significance levels varied, but at least one comparison was significant for most systems. Results were qualitatively similar in regression models adjusting for age (Fig. 3, Fig. S4): trends were almost universally toward higher PD levels predicting frailty, and effects were often but not always significant. Magnitudes of effects were relatively similar across systems, though the oxygen transport system tended to have slightly stronger effects, and the lipid system weaker effects. This was replicated with three types of regression: logistic with non-frail and prefrail pooled as the reference category (Fig. 3a); ordinal (Fig. 3b); and using the number of frailty criteria in a Poisson regression (Fig. 3c).

**Figure 2.**
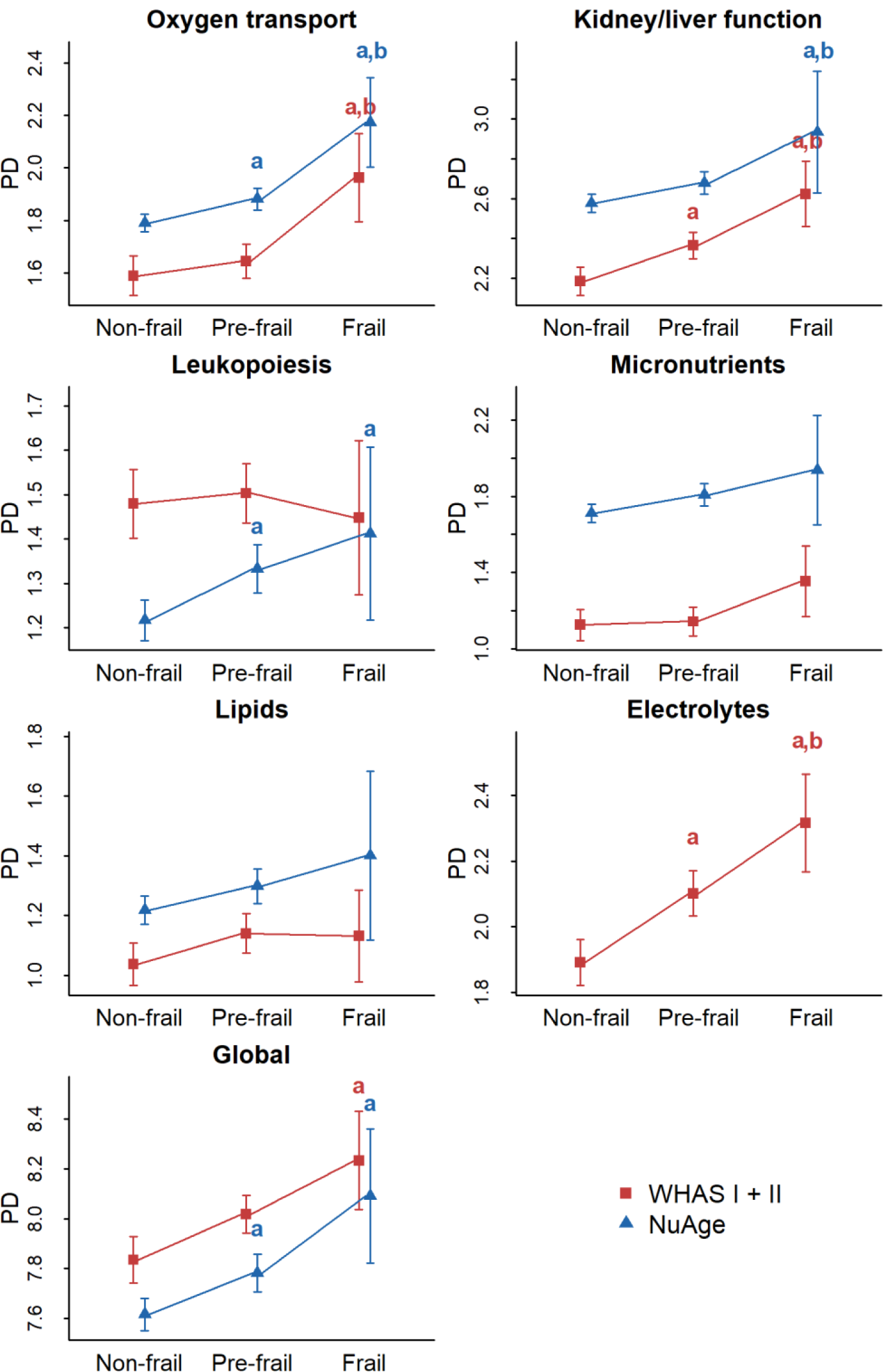
Physiological dysregulation levels by frailty phenotype for each system. Mean physiological dysregulation (PD) scores are shown in red for WHAS (I + II) and in blue for NuAge with corresponding 95% confidence intervals. Linear regressions were performed with frailty phenotype as a fixed effect, controlling for age with a cubic spline and for individual as a random effect when appropriate. Lower letter “a” indicates significantly different from the non-frail phenotype, while lower letter “b” indicates significantly different from the pre-frail phenotype.

**Figure 3.**
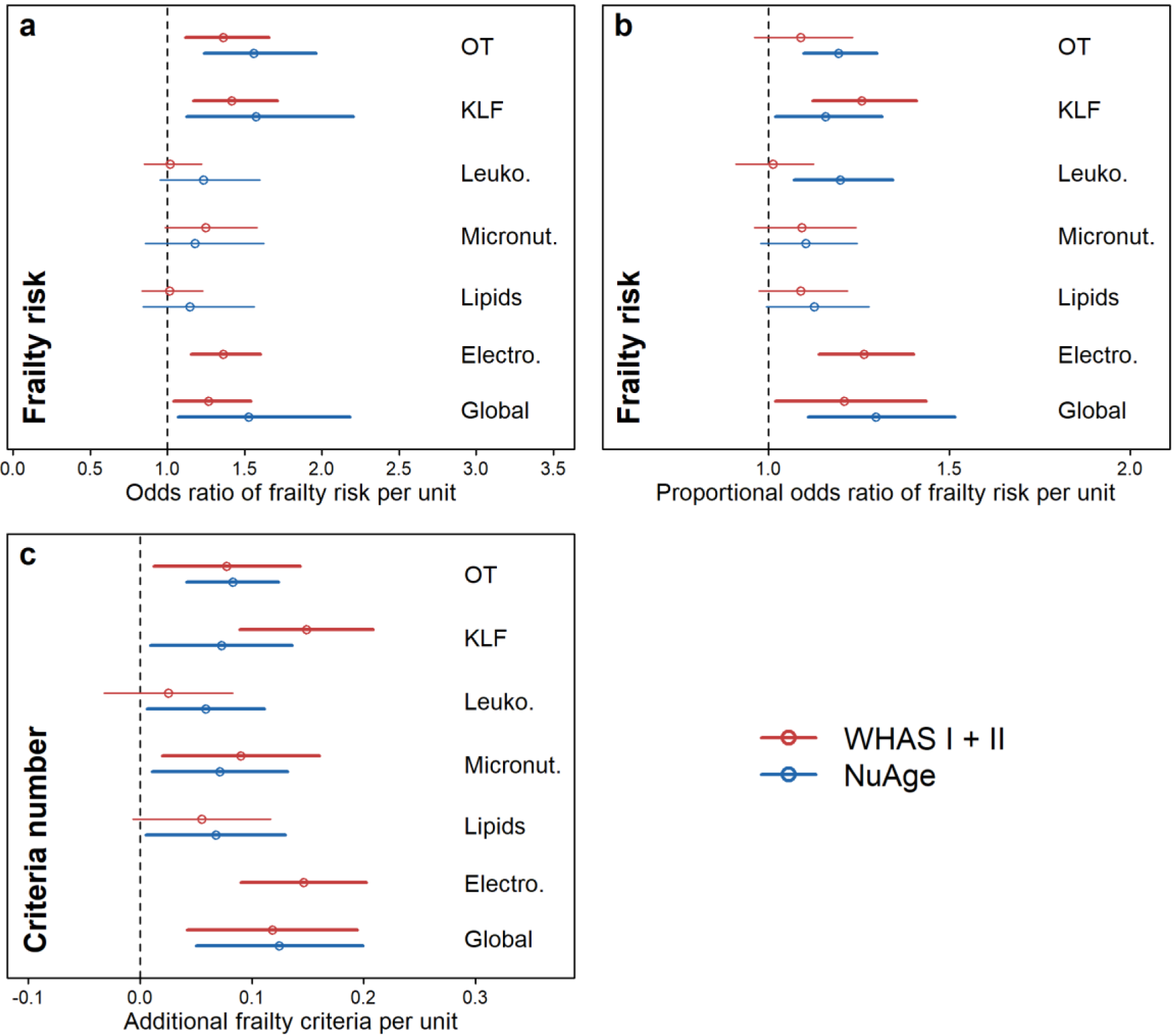
Association between physiological dysregulation and risk of frailty by system. Frailty risk was assessed through a logistic regression model with both non-frail and pre-frail in the reference group (**a**), through a proportional odds model (**b**), and with a Poisson regression using the number of frailty criteria (**c**). Estimations (points) together with 95% CIs (segments) are shown by specific-system and global dysregulation in red for WHAS (I + II) and in blue for NuAge. Abbreviations: Electro., Electrolytes; KLF, Kidney/liver function; Leuko., Leukopoiesis; Micronut., Micronutrients; OT, Oxygen transport.

### Association between physiological dysregulation and risk of individual frailty criteria

As expected, individual frailty criteria replicated the findings for global frailty, though more weakly (Fig. 4, Fig. S5). For most criteria, there was a trend toward greater prevalence with increasing dysregulation levels in the vast majority of analyses (dataset-system combinations), though significance levels varied. The exception was weakness, where there was no evidence of a trend across analyses and no analyses were significant. Overall, there were no dysregulation system-frailty criterion pairs that showed substantially stronger or weaker associations than the others, in the sense that confidence intervals were largely overlapping and any apparent differences in one dataset tended not to be replicated in the other.

**Figure 4.**
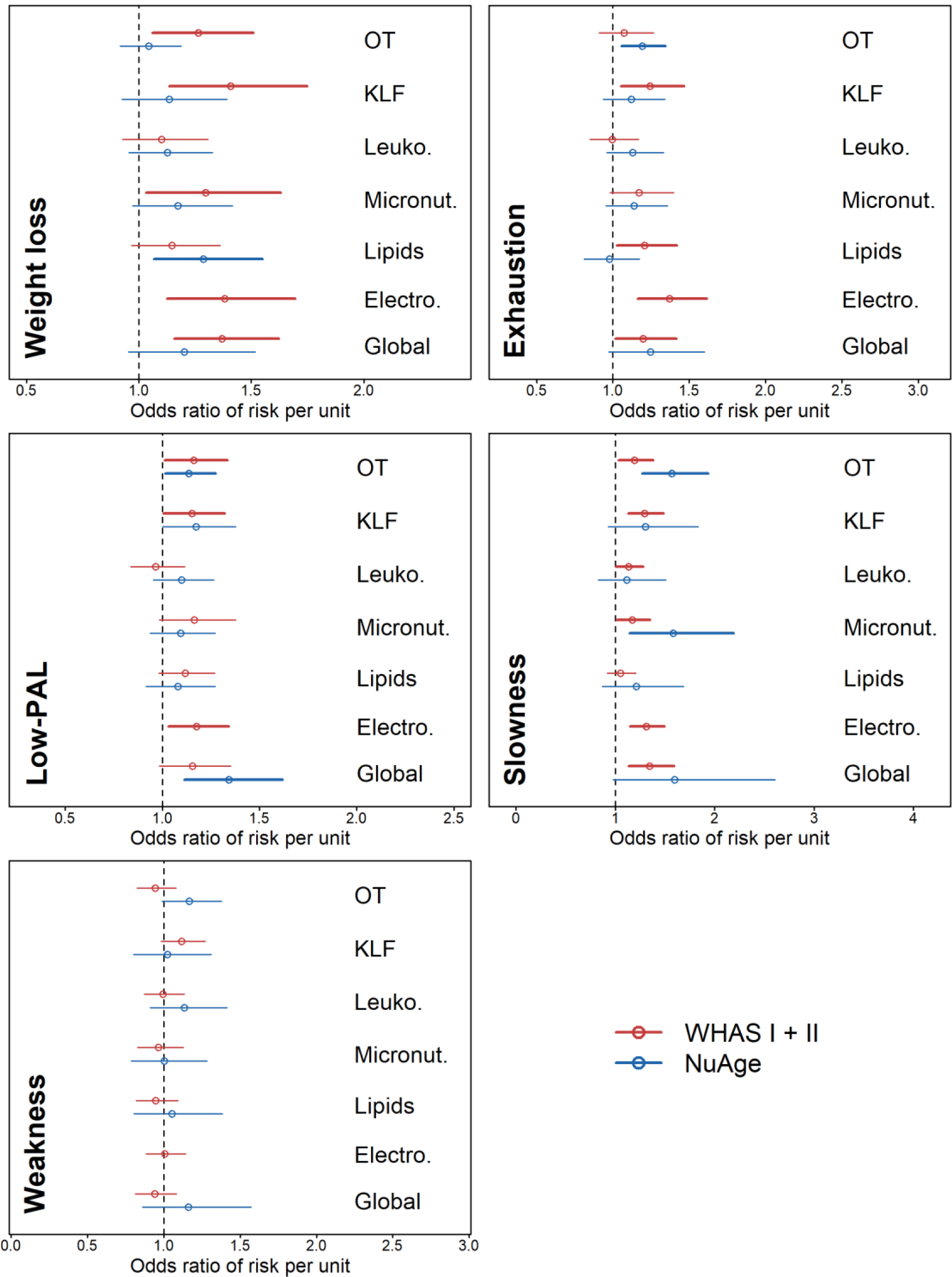
Association between physiological dysregulation and risk of individual frailty criteria. Estimations (points) together with 95% CIs (segments) for relationships between dysregulation levels and frailty risk (Odds ratio) are shown by specific-system and global dysregulation in red for WHAS (I + II) and in blue for NuAge. Abbreviations: Electro., Electrolytes; KLF, Kidney/liver function; Leuko., Leukopoiesis; Micronut., Micronutrients; OT, Oxygen transport.

### Linearity of associations between frailty and physiological dysregulation

Frailty status increased with the total cross-system PD, as represented by a weighted sum (Figs 5a and S6a). In both WHAS (I and II) and NuAge, the difference between frail and pre-frail was greater than between pre-frail and non-frail, suggesting accelerating frailty status with increasing PD, despite having already log-transformed the PD sum. Nonetheless, this nonlinearity was not statistically significant (Table S4); formal tests for non-linearity require greater statistical power, which is challenging with a small number of frail individuals. Likewise, the percent of frail individuals increased with number of dysregulated systems, and the percent of non-frail decreased (Figs 5b and S6b-c). This was true for both datasets, though more markedly so for WHAS. NuAge was underpowered to look at frailty by number of systems, so we included pre-frail with frail in this case. Figure S7 represents this differently, with all three frailty categories separated. Additionally, we tested for nonlinearity in individual physiological systems as their ability to predict frailty status in quadratic logistic regression models (Table S5). No significant quadratic effects were found in the full datasets.

**Figure 5.**
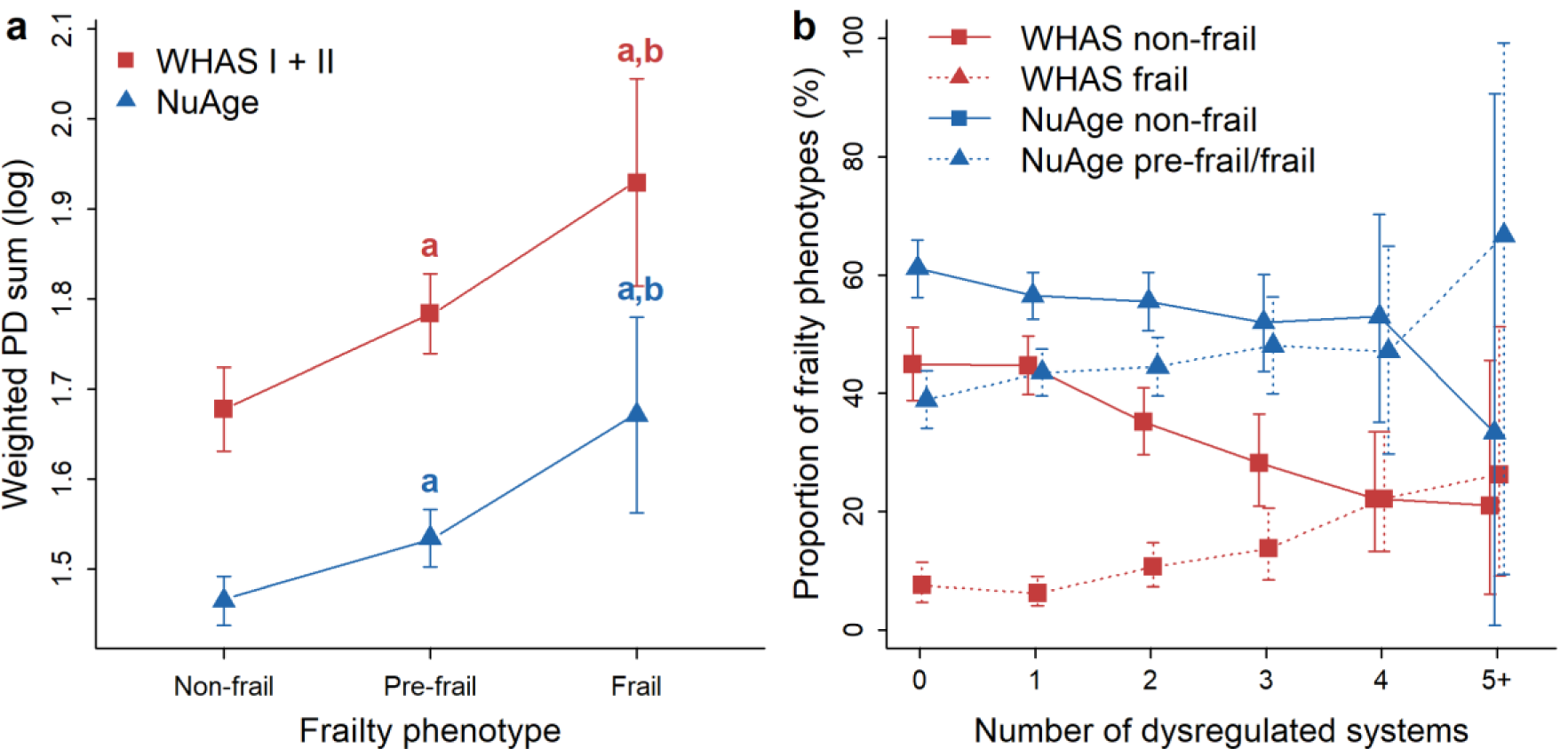
Cumulative effect of the number of dysregulated systems on frailty phenotype. **(a)** Mean levels of a weighted sum of PD across six (WHAS I + II, red) and five (NuAge, blue) systems by frailty phenotypes. Linear regressions were performed with frailty phenotype as a fixed effect, controlling for age with a cubic spline and for individual as a random effect. Lower letter “a” indicates significantly different from the non-frail phenotype, while lower letter “b” indicates significantly different from the pre-frail phenotype. **(b)** Prevalence of frail (dotted lines) and non-frail (solid lines) individuals according to the number of dysregulated systems in WHAS (red) and NuAge (blue) cohorts. Because the frail phenotype represents only 3% of all observations in NuAge, we combined pre-frail and frail subjects for this cohort.

### Association between the number of dysregulated physiological systems and risk of frailty

Independently of age, the risk to be frail significantly increased with the number of dysregulated physiological systems in both datasets (Fig. 6a grouping pre-frail with non-frail, Fig. S8a grouping pre-frail with frail; see also Fig. S9). There was a dose–response relationship between the number of dysregulated systems and risk of frailty that was clearly replicated across datasets and different ways to group the number of systems.

**Figure 6.**
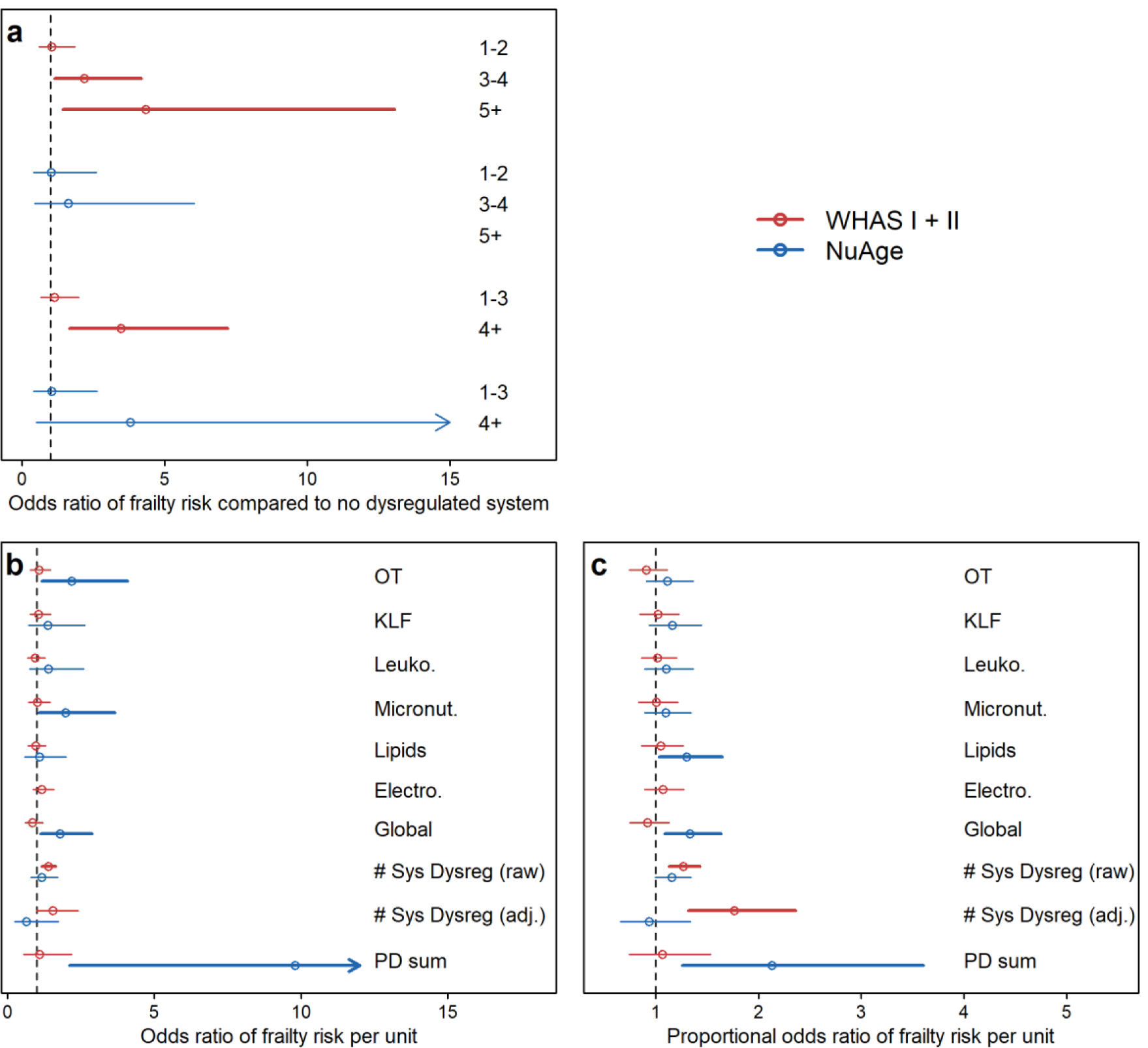
Association between physiological dysregulation and frailty risk in full regression models. Estimations (points) together with 95% CIs (segments) are plotted for relationships between dysregulation levels, or the number of dysregulated systems (“# Sys Dysreg”), and frailty risk in WHAS (n = 1194, red) and NuAge (n = 1653, blue). **(a)** Frailty risk was assessed with a logistic regression model comparing non-frail/pre-frail to frail (see Fig. S8 for models comparing non-frail to pre-frail/frail), as a function of the number of dysregulated systems categorized either as 1-2, 3-4, and 5+ (upper part) or 1-3 and 4+ (lower part), with no dysregulated system as the reference group. **(b)** Frailty risk was assessed with a logistic regression model comparing non-frail and pre-frail to frail, as a function of the number of dysregulated systems. **(c)** Frailty risk was also assessed with a proportional odds model on non-frail to pre-frail to frail. All models except “# Sys Dysreg (raw)” controlled for the presence or absence of dysregulation in individual systems (dummy variables). Abbreviations: # Sys Dysreg, Number of dysregulated systems; Electro., Electrolytes; KLF, Kidney/liver function; Leuko., Leukopoiesis; Micronut., Micronutrients; OT, Oxygen transport.

In WHAS, none of the individual systems significantly predicted frailty status after controlling for the number of systems, either relative to non-frail + pre-frail (Fig. 6b), or stepwise in an ordinal regression (Fig. 6c). However, the number of systems did significantly predict frailty status, and these effects actually grew stronger, though with larger confidence intervals, after controlling for which systems were dysregulated (Fig. 6, Fig. S8). On the other hand, the continuous cross-system measure of PD, PD sum, was not significant after control for which systems were dysregulated in WHAS. NuAge showed the opposite pattern (Fig. 6, Fig. S8): lipid and global dysregulation were significant after control for number of systems, but number of systems was not a significant predictor of frailty. In contrast, the PD sum measure was the best predictor of frailty status, even after control for the individual systems. Note, however, that confidence intervals overlap between the datasets, so this discrepancy is not necessarily an indication of qualitative differences.

## Discussion

Here, we tested a series of predictions that frailty is an emergent property of complex systems dynamics and breakdown in homeostasis of multiple physiological systems with age, namely *breadth* (frailty is associated with dysregulation of a wide variety of systems), *diffuseness* (frailty is associated with a variety of dysregulation profiles, with no specific system being necessary; analogous to distributed control in engineered systems (Guan et al. 2010), and *nonlinearity* (frailty emerges via a threshold effect, or risk accelerates with increasing cross-system dysregulation). Our results broadly but not universally supported these predictions.

Evidence for breadth was quite strong, with both datasets showing relatively similar effects of different systems on frailty risk, and no individual systems seeming particularly important. These results, using a data-driven rather than theory-driven set of systems and an additional dataset, replicate the study of Fried et al. (2009), which analyzed eight systems identified via abnormal biomarker levels in WHAS. They found no system that alone had a predictive value for frailty greater than 3.6-fold. Likewise, Li et al. (2015) found that five of six systems significantly predicted number of frailty criteria in at least one of two datasets (WHAS or InCHIANTI), but with no system markedly better than others. Lastly, Cappola et al. (2009) also found that three anabolic hormonal deficiencies had relatively similar augmented prevalences among frail individuals. More broadly, frailty has been associated with changes in a wide range of physiological systems (e.g. heart rate variability changes (Varadhan et al. 2009), oxidative stress (El Assar et al. 2020), changes in energy metabolism hormones (Kalyani et al. 2012), inflammation (Soysal et al. 2016) and inflamm-aging (Bandeen-Roche et al. 2009; Morrisette-Thomas et al. 2014), cortisol cycles (Varadhan et al. 2008), integrated albunemia (Cohen et al. 2015b), glucose/insulin signaling (Kalyani et al. 2012), and skeletal muscle ATP kinetics (Akki et al. 2014)).

Evidence for diffuseness is more nuanced. In no case were the odds ratios for a single system to predict frailty phenomenally high (e.g. > 10) as would be expected if a single system were key; such strong predictions only emerge when looking jointly at the number of systems, as we show here or in Fried et al. (2009). Strong predictions of frailty by number of dysregulated systems with no residual effects of which systems were dysregulated would also provide clear evidence of diffuseness. This was the case in WHAS, with the minor nuance that one adjusted model was not quite statistically significant, even though the effect size increased from the significant unadjusted model. However, the sum of dysregulation was not associated with frailty risk after adjustment for which systems were dysregulated. Conversely, in NuAge, most but not all systems failed to predict frailty after adjustment for number of systems, but number of systems also failed to predict frailty, and only the sum of PD was predictive. We believe the most likely explanation for this discrepancy is the more general difficulty of replicating subtle effects without overwhelming statistical power, particularly in this context of NuAge being healthier, and with slightly different definitions of frailty criteria. There was not a qualitative difference between results from the two datasets (confidence intervals overlapping). To our knowledge, the only evidence of diffuseness in the literature is from Fried et al. (2009), who showed that number of dysregulated systems remained significant after adjustment for which systems, and only one of eight systems remained significant after adjusting for the number.

Evidence for nonlinearity was also ambiguous, largely due to limited sample size of frail individuals and individuals with many systems dysregulated. Individual systems did not mostly show significant nonlinear associations with frailty risk. There was a clear trend for acceleration of frailty risk with greater sum of PD in both datasets, though this was not significant. For the number of systems, the trend is visually nonlinear in WHAS, both for the increase in frailty and the decrease in non-frail. For NuAge, the small sample sizes make it hard to discern. Fried et al. (2009) showed somewhat more convincing evidence for nonlinearity.

Our findings are broadly but not perfectly consistent with those of Fried et al. (2009). The major discrepancy is that Fried et al. (2009) provided clearer evidence of nonlinearity and diffuseness. We note, however, that an apples-to-apples comparison (restricting ourselves to WHAS and to the number of systems rather than the PD sum measure), our findings do replicate Fried et al. (2009) in this separate set of systems. The cases where the broader replication fails are cases where our results are ambiguous (i.e., underpowered), rather than contradictory. We thus believe that these findings show that a similar systems model of frailty can be derived from quite distinct definitions of physiological systems, and that this is in fact one of the stronger arguments for viewing frailty as an emergent phenomenon of the complex systems dynamics.

Overall, our analyses thus show a mix of results supporting the predictions of frailty as an emergent property of multisystem dysregulation, and results where we are underpowered to draw conclusions one way or the other. This is exacerbated by measurement error, which is likely to be quite large for both frailty itself and for physiological dysregulation. This is particularly problematic for the criterion of nonlinearity: as measurement error of the two variables increases, a threshold effect will become a sigmoidal curve and subsequently a linear relationship. More generally, in many studies, the representation of frail individuals is relatively low, as is the representation of individuals with many systems dysregulated. Further confirmation from other studies will be required, or perhaps a meta-analysis that would increase power relative to individual studies while avoiding the problems that could arise from data harmonization.

Even if we were to accept that the current evidence were sufficient to prove breadth, diffuseness, and nonlinearity, how good would that evidence be for frailty as an emergent property of a complex system? On the one hand, it would be insufficient to rule out alternative hypotheses. For example, perhaps the various systems measured here and in other studies all show age-related and health-related deterioration. If frailty tends to correlate with health state in general, it would also correlate with a series of individual systems and indicators that correlate strongly with general health. Both breadth and diffuseness could easily be satisfied, and nonlinearity as well if patterns of health decline are broadly exponential. While we cannot rule out such an explanation, it would require that frailty have another cause. It is not impossible that some as-yet-undiscovered physiological pathway underlies frailty, or perhaps some pathway that no one has thought to analyze in this context. However, it seems unlikely that such a big gap in our knowledge of physiological changes with age would exist at this point. We consider such a scenario unlikely but not impossible.

On the other hand, the scenario presented here is not the only possibility for how frailty might emerge from complex systems dynamics. For example, frailty might be the result of a critical transition from a stable physiological state, or might itself be the critical transition to death (Nakazato et al.; Gijzel et al. 2017). In order to distinguish between these possibilities, longitudinal data on a fine temporal scale will be required. Alternatively, frailty might represent a loss of redundancy in physiological networks leading to declining robustness (Gavrilov and Gavrilova 2004; Kitano 2004; Nijhout et al. 2017), or might actually represent an adaptation to other changing conditions (Cohen et al. in press; Le Couteur and Simpson 2011). Nearly all of the evidence presented here – both from this study and the literature – is cross-sectional, making it hard to disentangle such possibilities, which are not mutually exclusive.

In addition to the important caveats above, we add a few other limitations. First, the six systems measured here are a small fraction of the systems that might be measured, and were not chosen based on any *a priori* reason to expect them to be associated strongly with frailty. We note, however, the strong concordance between our results and those of Fried et al. (2009), with two very difference sets of systems. Second, the systems here are limited to those that can be measured through circulating clinical biomarkers, and may not be representative of other organs and tissues. Third, we are using the phenotypic definition of frailty, which is somewhat hard to standardize across studies due to differing measurement instruments and percentile cut-offs (Theou et al. 2015). Accordingly, the comparison between NuAge and WHAS is imprecise. Fourth, for the prediction of nonlinearity, it is difficult to distinguish a prediction between a sigmoidal relationship (likely if there is an underlying physiological threshold beyond which frailty is reached) and an exponential relationship (frailty risk increases indefinitely as dysregulation increases). Empirically, if the threshold is toward the upper range of physiological dysregulation observed and is slightly variable, the two scenarios may be indistinguishable. We note that if we were able to detect a true threshold effect, we should expect 100% prevalence beyond the threshold. Fifth, some of the biomarkers, notably lipids and micronutrients, can be influenced by medication (e.g. statins) and supplements.

Despite these caveats, this study contributes to a growing body of literature suggesting that phenotypic frailty is an emergent property of the complex systems dynamics as physiological regulation breaks down toward the end of life. It is also consistent with the loosely related construct of the frailty index (Mitnitski et al. 2001; Rockwood et al. 2005). Phenotypic frailty and the frailty index, despite their names, describe distinct phenomena; nonetheless, the frailty index as well shows some properties of a complex system, and has been linked to network dynamics (Mitnitski et al. 2017; Rutenberg et al. 2018). Future research should aim to increase power, either with larger studies or via meta-analysis, in order to confirm whether the diffuseness and nonlinearity are indeed generally present. It will also be critical to assess whether frailty is an outcome of a critical transition, i.e. an alternative stable state, or is itself the process of the critical transition toward death. In the former case, rates of change in underlying physiological indices should slow and stabilize during frailty; in the latter, they should increase. Fine scale time-series data will be necessary.

Lastly, it will be important to establish to what extent frailty is a global physiological process. At this point, it seems increasingly unlikely that frailty is a highly local process (i.e., a product of one or two specific systems), but this does not imply that it is fully global. Are there some systems that do not change during frailty? Does the vulnerability to stressors apply uniformly to all types of stress, or to specific stressors? Given our current state of knowledge, frailty as an emergent state based on underlying physiological dynamics seems to be the most probable explanation, but many questions remain.

## Declarations

## Ethics approval

Both studies were approved by their local Ethic Committee, in terms of study protocol and consent procedure. The NuAge Database and Biobank and secondary analysis for this project were approved by the *Comité d’éthique de la recherche du CIUSSS de l’Estrie – CHUS* (projects 2019-2832 and 14-059, respectively). All participants recruited in both studies signed informed consent.

## Code availability

All code is available upon request.

