## Supplementary material for "The frailty syndrome as an emergent state of parallel dysregulation in multiple physiological systems": Table S1, Table S2, Table S3, Table S4, Table S5,Fig S1, Fig S2, Fig S3, Fig S4, Fig S5, Fig S6, Fig S7, Fig S8, Fig S9

### Supplementary Tables

**Table S1.** Summary statistics of biomarkers at baseline, indicated as mean ± standard deviation.

| Biomarker | System | WHAS | | | NuAge | | |
|---|---|---|---|---|---|---|---|
| | | I | II | I + II | All | Men | Women |
| Ferritin (μg/L)[a] | OT | 119.1 ± 129.9 | 111.7 ± 99.4 | 115.3 ± 120.0 | – | – | – |
| Hemoglobin (g/L) | OT | 129.5 ± 13.9 | 133.0 ± 11.3 | 131.8 ± 12.6 | 138.9 ± 12.6 | 145.0 ± 12.0 | 133.3 ± 10.3 |
| Hematocrit (%) | OT | 39.6 ± 4.2 | 40.0 ± 3.3 | 39.9 ± 3.7 | 41.1 ± 3.7 | 42.8 ± 3.5 | 39.5 ± 3.0 |
| Iron (μmol/L) | OT | 13.7 ± 4.6 | 14.7 ± 4.9 | 14.4 ± 4.9 | – | – | – |
| Mean corpuscular hemoglobin (MCH, pg) | OT | 30.5 ± 2.3 | 30.6 ± 2.0 | 30.5 ± 2.2 | 31.0 ± 1.7 | 31.3 ± 1.6 | 30.8 ± 1.7 |
| MCH concentration (g/L) | OT | 327.7 ± 13.3 | 332.6 ±13.7 | 330.6 ± 13.2 | 338.5 ± 7.6 | 339.4 ± 7.3 | 337.8 ± 7.8 |
| Red blood cells ($10^{12}$/L) | OT | 4.27 ± 0.48 | 4.35 ± 0.36 | 4.33 ± 0.40 | 4.49 ± 0.42 | 4.64 ± 0.42 | 4.34 ± 0.37 |
| Red cell distribution width (%)[a] | OT | – | – | – | 13.5 ± 0.9 | 13.6 ± 0.8 | 13.5 ± 1.0 |
| Albumin (g/L) | KLF | 40.4 ± 3.0 | 42.6 ± 2.7 | 41.7 ± 3.2 | 39.2 ± 2.9 | 39.5 ±2.9 | 38.9 ± 2.9 |
| Alkaline phosphatase (U/L)[a] | KLF | 96.6 ± 44.0 | 84.8 ± 26.6 | 90.1 ± 36.2 | 78.2 ± 23.2 | 76.2 ± 22.9 | 80.1 ± 23.5 |
| Alanine transaminase (U/L)[a] | KLF | 15.8 ± 14.2 | 16.8 ± 7.4 | 16.5 ± 8.9 | 11.4 ± 5.4 | 12.5 ± 5.7 | 10.3 ± 4.9 |
| Aspartate transaminase (U/L)[a] | KLF | 18.8 ± 11.2 | 19.4 ± 6.5 | 19.0 ± 7.0 | 21.7 ± 5.4 | 22.6 ± 5.9 | 20.8 ± 4.8 |
| Gamma-glutamyl transferase (U/l)[a] | KLF | 32.9 ± 45.0 | 27.6 ± 28.4 | 29.8 ± 33.2 | 32.7 ± 28.6 | 35.9 ± 25.9 | 29.4 ± 30.8 |
| Protein, total (g/L) | KLF | 69.0 ± 5.0 | 71.1 ± 4.6 | 70.2 ± 4.8 | 72.4 ± 4.1 | 72.6 ± 4.0 | 72.2 ± 4.1 |
| Uric acid (μmol/L) | KLF | 352.4 ± 105.4 | 319.2 ± 86.2 | 331.2 ± 93.4 | 338.0 ± 79.2 | 365.4 ± 74.2 | 310.3 ± 74.5 |
| Basophils (%)[b] | Leuko. | 0.77 ± 0.50 | 0.72 ± 0.52 | 0.73 ± 0.48 | 0.15 ± 0.44 | 0.14 ± 0.42 | 0.15 ± 0.46 |
| Lymphocytes (%) | Leuko. | 29.3 ± 9.7 | 29.1 ±8.2 | 29.2 ± 8.7 | 29.1 ± 7.7 | 27.7 ± 7.4 | 30.4 ± 7.8 |
| Monocytes (%) | Leuko. | 7.05 ± 2.47 | 6.90 ± 2.36 | 7.00 ± 2.37 | 7.21 ± 2.64 | 7.49 ± 2.84 | 6.96 ± 2.42 |
| Neutrophils (%) | Leuko. | 60.3 ± 10.2 | 60.1 ± 10.0 | 60.2 ± 9.8 | 63.5 ± 8.0 | 64.6 ± 8.0 | 62.4 ± 7.9 |
| White blood cells ($10^9$/L)[a] | Leuko. | 6.58 ± 3.81 | 6.27 ± 1.67 | 6.38 ± 1.76 | 6.16 ± 1.59 | 6.25 ± 1.59 | 6.07 ± 1.58 |

| | | | | | | | |
|---|---|---|---|---|---|---|---|
| 25-hydroxy vitamin D (nmol/L)[a] | Micronut. | 52.2 ± 26.3 | 55.1 ± 24.5 | 54.8 ± 26.2 | – | – | – |
| Folate (nmol/L)[a] | Micronut. | 26.9 ± 31.1 | 27.1 ± 19.4 | 27.7 ± 30.5 | 45.0 ± 11.0 | 45.1 ± 11.1 | 45.0 ± 11.0 |
| Vitamin A (µmol/L) | Micronut. | 2.62 ± 0.88 | 2.35 ± 0.73 | 2.48 ± 0.83 | – | – | – |
| Vitamin B12 (pmol/L)[a] | Micronut. | 359.4 ± 192.7 | 377.2 ± 305.6 | 372.6 ± 263.7 | 354.2 ± 141.3 | 339.2 ± 134.5 | 369.6 ± 146.6 |
| α-Tocopherol (µmol/L)[a] | Micronut. | – | – | – | 28.6 ± 13.2 | 26.8 ± 13.2 | 30.4 ± 12.9 |
| β-Carotene (µmol/L)[a] | Micronut. | – | – | – | 3.42 ± 6.84 | 3.05 ± 8.18 | 3.79 ± 5.14 |
| γ-Tocopherol (µmol/L)[a] | Micronut. | – | – | – | 3.27 ± 2.43 | 3.35 ± 2.59 | 3.20 ± 2.26 |
| Cholesterol, total (mmol/L) | Lipids | 5.81 ± 1.08 | 6.03 ± 1.00 | 5.97 ± 1.00 | 5.23 ± 0.97 | 4.96 ± 0.91 | 5.49 ± 0.96 |
| Cholesterol/HDL ratio | Lipids | 4.51 ± 1.43 | 4.46 ± 1.45 | 4.51 ± 1.42 | 3.88 ± 1.05 | 4.12 ± 1.09 | 3.64 ± 0.95 |
| High-density lipoproteins (mmol/L) | Lipids | 1.38 ± 0.39 | 1.47 ± 0.43 | 1.43 ± 0.41 | 1.42 ± 0.39 | 1.27 ± 0.38 | 1.58 ± 0.38 |
| Low-density lipoproteins (mmol/L) | Lipids | – | – | – | 3.07 ± 0.82 | 2.96 ± 0.79 | 3.18 ± 0.84 |
| Triglycerides (mmol/L)[a] | Lipids | 1.97 ± 1.20 | 1.73 ± 1.00 | 1.87 ± 1.14 | 1.65 ± 0.77 | 1.65 ± 0.79 | 1.66 ± 0.75 |
| Calcium (mmol/L) | Electro. | 2.32 ± 0.11 | 2.38 ± 0.10 | 2.36 ± 0.11 | – | – | – |
| Chloride (mmol/L) | Electro. | 102.4 ± 3.9 | 102.7 ± 3.2 | 102.6 ± 3.6 | – | – | – |
| Potassium (mmol/L) | Electro. | 4.14 ± 0.43 | 4.14 ±0.37 | 4.12 ± 0.39 | – | – | – |
| Magnesium (mmol/L) | Electro. | 0.82 ± 0.09 | 0.83 ± 0.08 | 0.82 ± 0.08 | – | – | – |
| Phosphate (mmol/L) | Electro. | 1.17 ± 0.17 | 1.16 ± 0.16 | 1.17 ± 0.17 | – | – | – |
| Sodium (mmol/L) | Electro. | 139.0 ± 3.2 | 139.7 ± 2.5 | 139.4 ± 3.0 | – | – | – |

Note: Lower letter "a" indicates variables that were log-transformed, while lower letter "b" indicates variables that were square root-transformed. Abbreviations: Electro., Electrolytes; KLF, Kidney/Liver function; Leuko., Leukopoiesis; Micronut., Micronutrients; MCH, Mean corpuscular hemoglobin; OT, Oxygen transport.

**Table S2.** Sample size (number of observations across all time points) per dataset and physiological system

| System | WHAS | | | NuAge | | |
|---|---|---|---|---|---|---|
| | I | II | I + II | All | Men | Women |
| Oxygen transport | 1508 | 1038 | 1707 | 5566 | 2684 | 2882 |
| Kidney/liver function | 1590 | 1081 | 1780 | 2985 | 1504 | 1481 |
| Leukopoiesis | 1456 | 978 | 1620 | 3255 | 1541 | 1714 |
| Micronutrients | 1439 | 734 | 1368 | 2849 | 1430 | 1419 |
| Lipids | 1602 | 1082 | 1788 | 2881 | 1434 | 1447 |
| Electrolytes | 1577 | 1047 | 1739 | – | | |
| Global | 1277 | 637 | 1196 | 1653 | 795 | 858 |

**Table S3.** Frailty criteria definitions

| | WHAS | NuAge |
|---|---|---|
| **Weight loss** | Unintentional weight loss ≥10 pounds in prior year or, at follow-up, or ≥5% of body weight in prior year (by direct measurement of weight). | Weight loss ≥ 5% (based on self-reported weight loss in the last 12 months at baseline, and by direct measurement of weight at follow-ups) or BMI < 22. |
| **Weakness** | Grip strength in the lowest 20% at baseline, adjusted for BMI. | Grip strength < (mean − 1 SD by age group (67 to 72, 73 to 77, and 78 to 84 years) and sex within NuAge. Grip strength was calculated as the maximum value of six assays (three for each hand), re-calculated at each follow-up. |
| **Exhaustion** | Self-reported exhaustion, identified by two questions from the CES–D scale. | Vitality subscale of the SF-36 (Ware & Sherbourne, 1992) score < (mean − 1 SD) by age group (67 to 72, 73 to 77, and 78 to 84 years) and sex within NuAge. |
| **Slowness** | The slowest 20% of the population was defined at baseline, based on time to walk 15 feet, adjusting for standing height. | Brisk walking pace < 1 m/s (Studenski et al., 2003). |
| **Low-PAL** | A weighted score of kilocalories expended per week calculated at baseline, based on each participant's report. The lowest quintile of physical activity was identified. | PASE (Washburn et al., 1993) score < (mean − 1 SD) by age group (67 to 72, 73 to 77, and 78 to 84 years) and sex within NuAge. |

Abbreviations: BMI, body mass index; CES-D, Center for Epidemiologic Studies Depression Scale; Low-PAL, low physical activity level; PASE, Physical Activity Scale for the Elderly; SF-36, Short Form (36) Health Survey.

**Table S4**. Non-linearity of frailty associations with summed system-specific dysregulation

| | WHAS I + II | | NuAge | |
|---|---|---|---|---|
| | **Effect [95% CI]** | ***p*** | **Effect [95% CI]** | ***p*** |
| **Model 1: Categorical** | | | | |
| Pre-frail (vs. non-frail) | **0.084 [0.01,0.16]** | **0.03** | **0.043 [0.003,0.08]** | **0.03** |
| Frail (vs. non-frail) | **0.24 [0.1,0.38]** | **0.0007** | **0.163 [0.05,0.28]** | **0.007** |
| **Model 2: Quadratic** | | | | |
| Frailty (Linear 0 - 1 - 2) | 0.048 [-0.1,0.19] | 0.51 | 0.005 [-0.08,0.09] | 0.90 |
| Frailty quadratic term | 0.036 [-0.05,0.12] | 0.40 | 0.038 [-0.02,0.10] | 0.23 |
| **Model 3: Linear + Categorical** | | | | |
| Frailty (Linear 0 - 1 - 2) | **0.084 [0.01,0.16]** | **0.03** | **0.04366 [0.004,0.08]** | **0.03** |
| Frail (vs.pre- and non-frail) | 0.071 [-0.1,0.24] | 0.40 | 0.077 [-0.05,0.20] | 0.23 |

Notes: Regression models predicting the log-sum of PD in all systems. CI: Confidence interval. Models 2 and 3 code frailty as a numeric term with non-frail=0, pre-frail=1, and frail=2. All models control for age as a cubic basis spline, and NuAge models also control for sex; sex and age terms were never significant ($p > 0.1$). Bold indicates $p < 0.05$, and italics indicate $0.05 < p < 0.1$.

**Table S5**. Non-linearity associations of system-specific dysregulation with frailty risk

| **System** | **WHAS I + II** | | **NuAge** | |
|---|---|---|---|---|
| | **OR [95% CI]** | ***p*** | **OR [95% CI]** | ***p*** |
| Oxygen transport | *0.94 [0.88-1.00]* | *0.07* | *0.92 [0.84-1.00]* | *0.06* |
| Kidney/liver function | 0.97 [0.90-1.04] | 0.32 | 1.01 [0.96-1.05] | 0.49 |
| Leukopoiesis | 0.97 [0.87-1.09] | 0.64 | 0.98 [0.93-1.04] | 0.59 |
| Micronutrients | *1.09 [0.99-1.19]* | *0.09* | 1.03 [0.92-1.16] | 0.57 |
| Lipids | 0.99 [0.89-1.11] | 0.90 | 1.01 [0.97-1.04] | 0.75 |
| Electrolytes | 0.92 [0.82-1.03] | 0.13 | – | |
| Global | 0.95 [0.89-1.02] | 0.13 | 0.75 [0.52-1.09] | 0.13 |

Notes: Odds ratio (OR) for the quadratic terms of regression models predicting frailty (frail vs non-frail/pre-frail) as a function of PD (not log-transformed), controlling for age, sex (NuAge), and repeated measures. Bold indicates $p < 0.05$, and italics indicate $0.05 < p < 0.1$. Abbreviation: CI, Confidence interval.

### Supplementary Figures

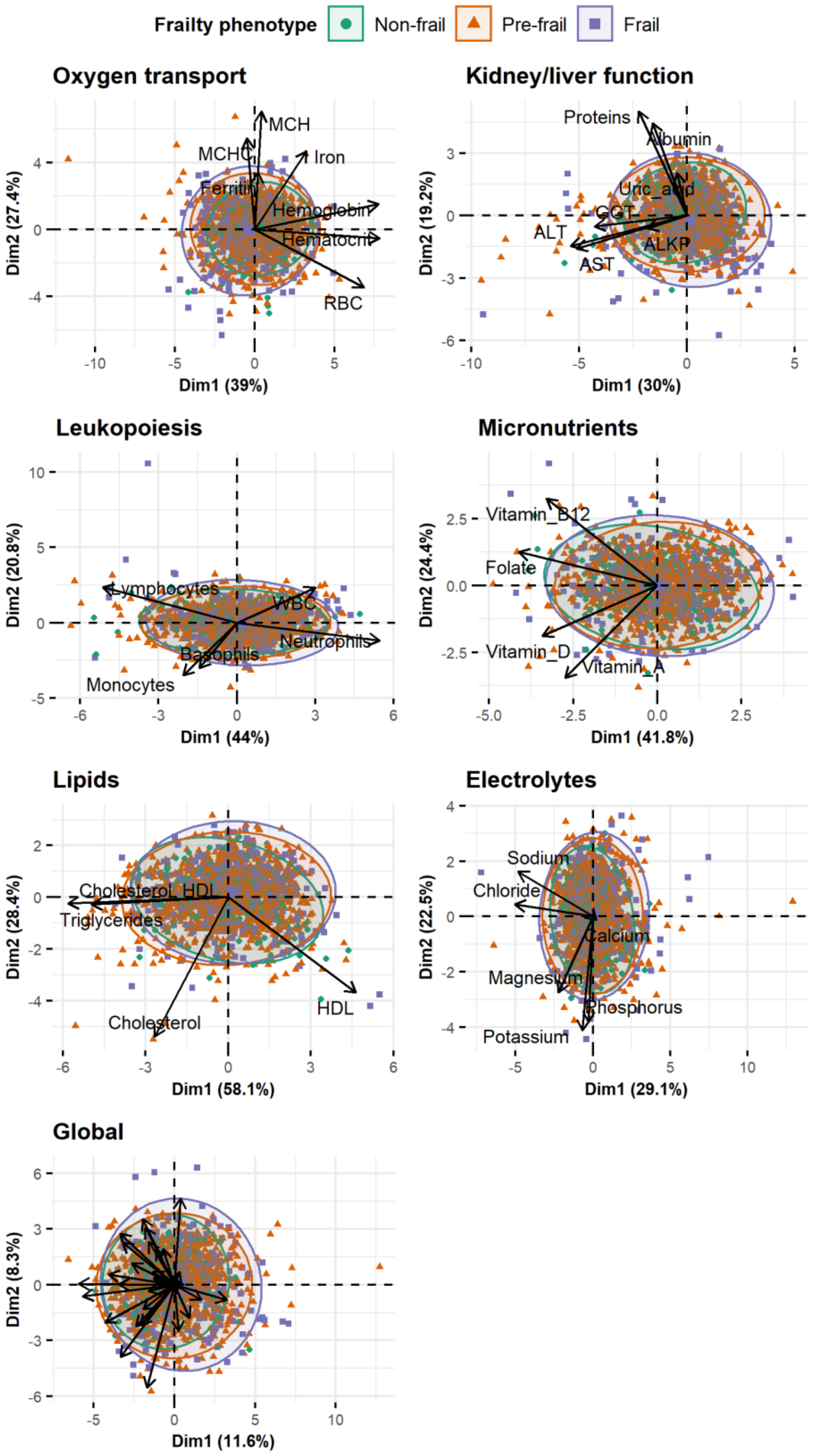


**Figure S1.** Principal component analyses to characterise biomarkers profile of non-frail *vs* pre-frail *vs* frail individuals in WHAS I for each physiological system and globally.

Individuals (points) together with 95% ellipses (dispersion) by frailty phenotypes for each physiological system are presented. Different colors indicate frailty phenotypes.

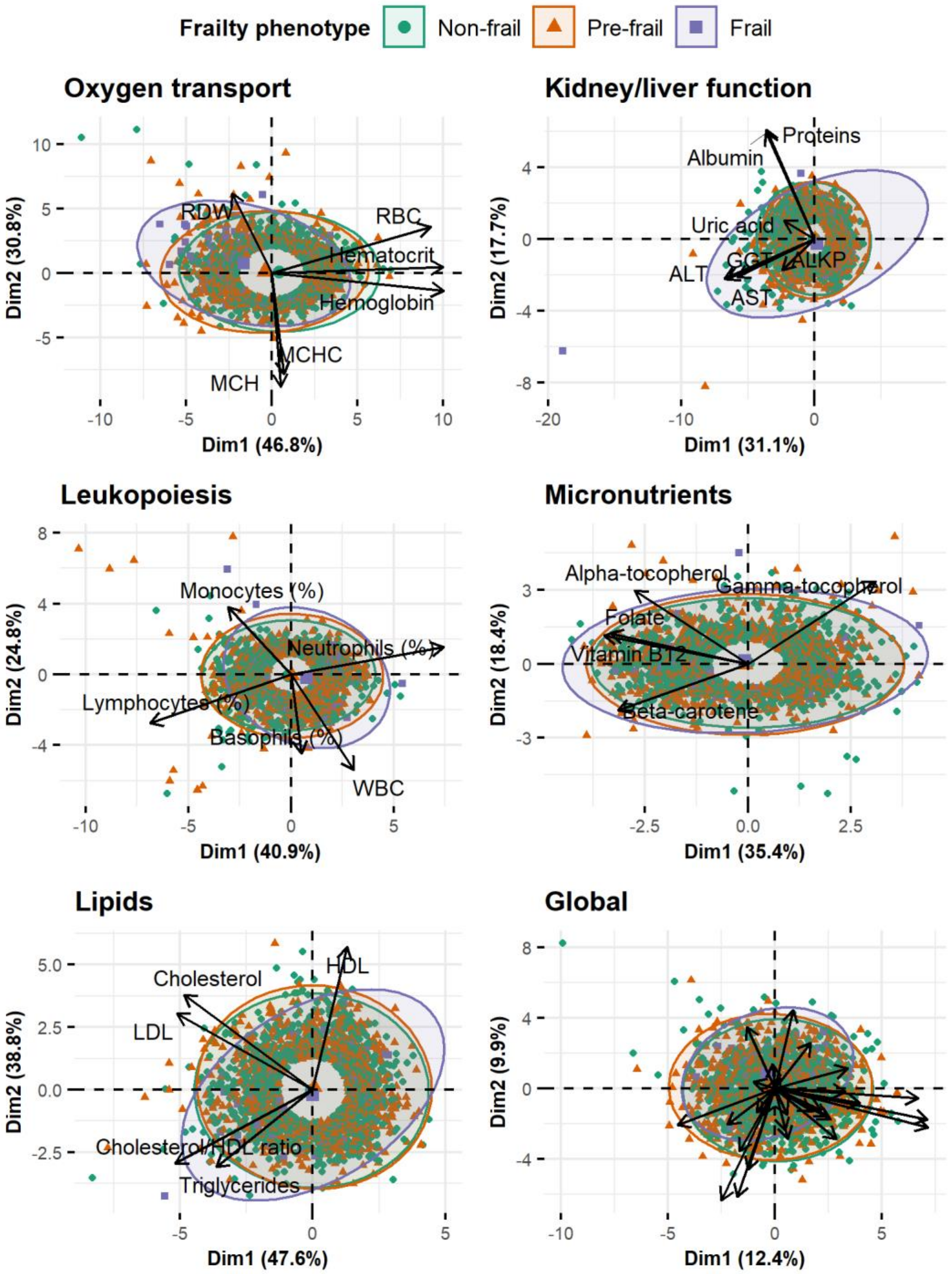


**Figure S2.** Principal component analyses to characterise biomarkers profile of non-frail *vs* pre-frail *vs* frail individuals in NuAge for each physiological system and globally. Individuals (points) together with 95% ellipses (dispersion) by frailty phenotypes for each physiological system are presented. Different colors indicate frailty phenotypes.

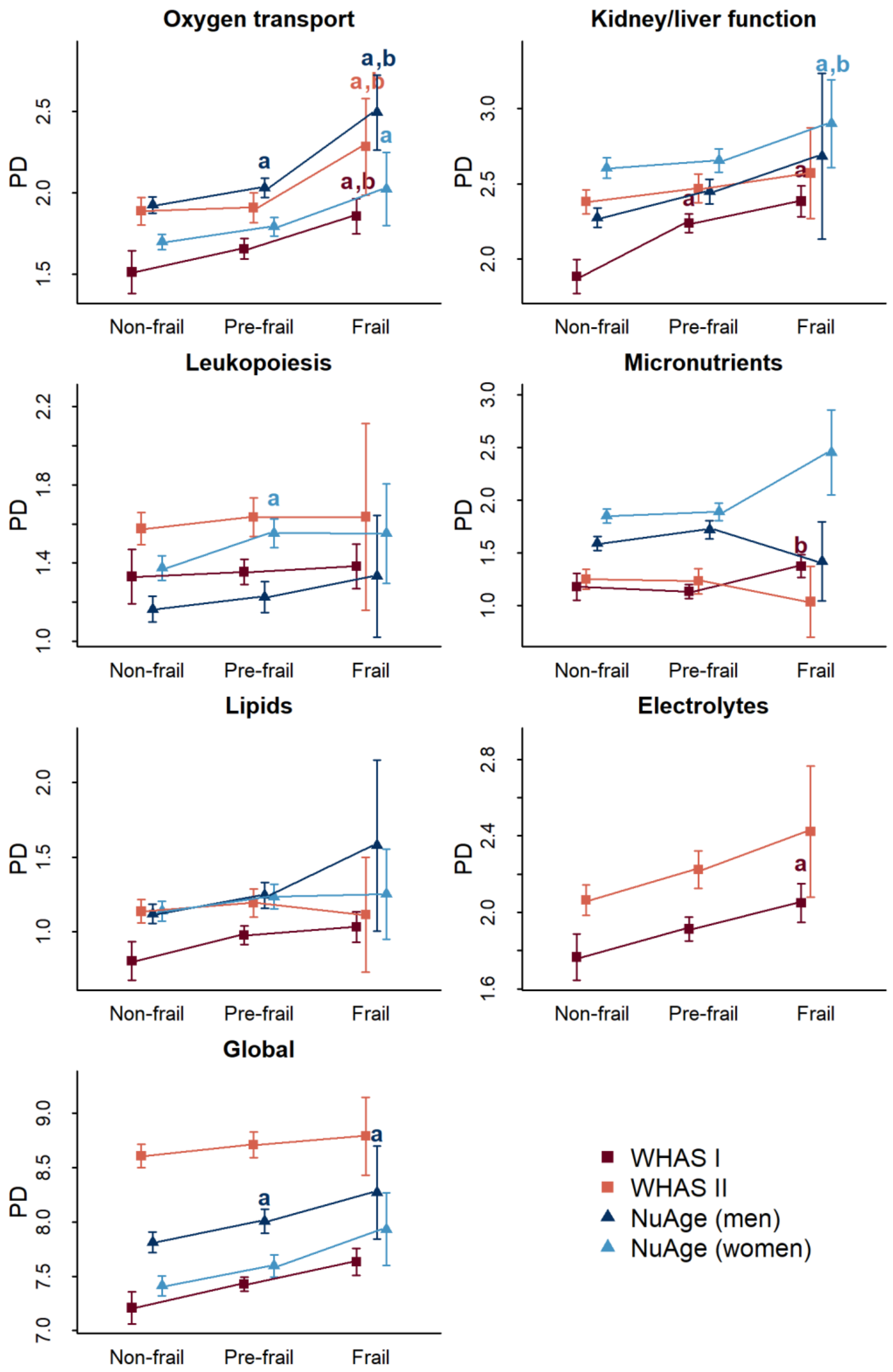


**Figure S3. Physiological dysregulation levels by frailty phenotype for each system in WHAS I, WHAS II, and by sex in the NuAge cohort**. Mean physiological dysregulation (PD) scores are shown with corresponding confidence intervals. Linear regressions were performed with frailty phenotype as a fixed effect, controlling for age with a cubic spline and for individual as a random effect when appropriate. “a” indicates significantly different from the non-frail phenotype, while “b” indicates significantly different from the pre-frail phenotype.

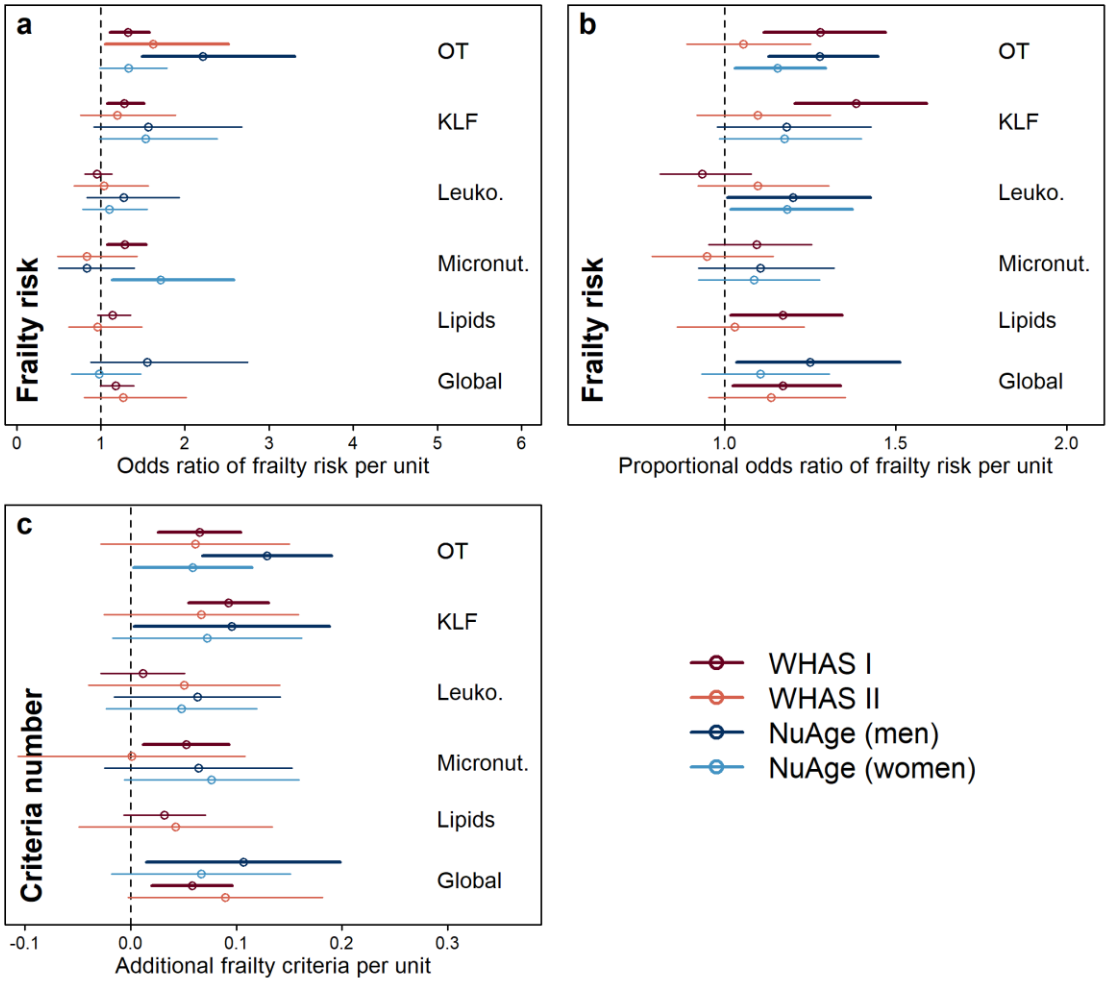


**Figure S4. Association between physiological dysregulation and risk of frailty by system in WHAS I and WHAS II, and by sex in the NuAge cohort**. Frailty risk was assessed through a logistic regression model with both non-frail and pre-frail in the reference group (**a**), through a proportional odds model (**b**), and with a Poisson regression using the number of frailty criteria (**c**). Estimations (points) together with 95% CIs (segments) are shown by specific-system and global dysregulation. Different colors indicate different datasets or (subsets). Abbreviations: Electro., Electrolytes; KLF, Kidney/liver function; Leuko., Leukopoiesis; Micronut., Micronutrients; OT, Oxygen transport.

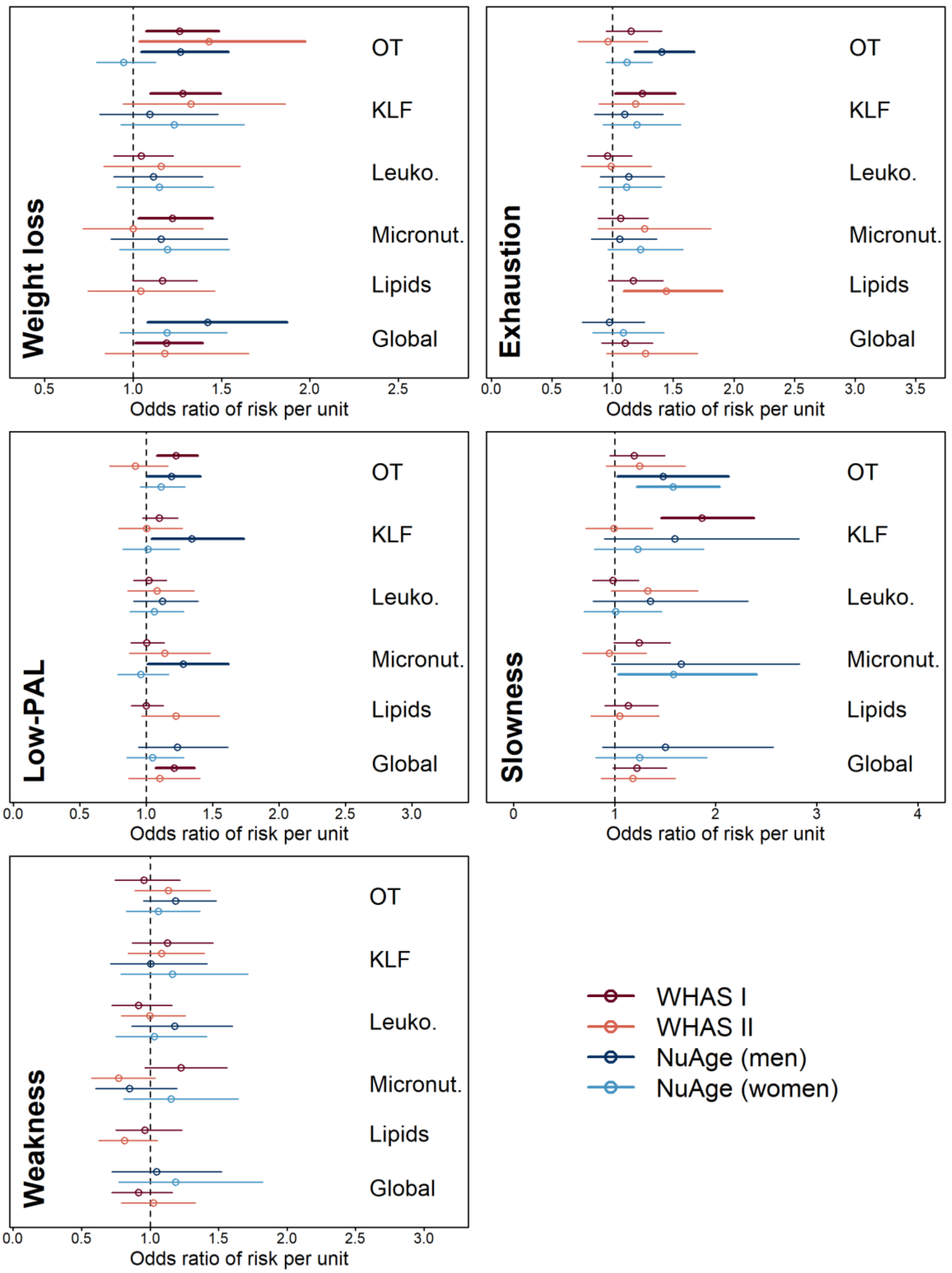


**Figure S5. Association between physiological dysregulation and risk of individual frailty criteria in WHAS I and WHAS II, and by sex in the NuAge cohort**. Estimations (points) together with 95% CIs (segments) for relationships between dysregulation levels and frailty risk (Odds ratio) are shown by specific-system and global dysregulation. Different colors indicate different datasets (or subsets). Abbreviations: KLF, Kidney/liver function; Leuko., Leukopoiesis; Micronut., Micronutrients; OT, Oxygen transport.

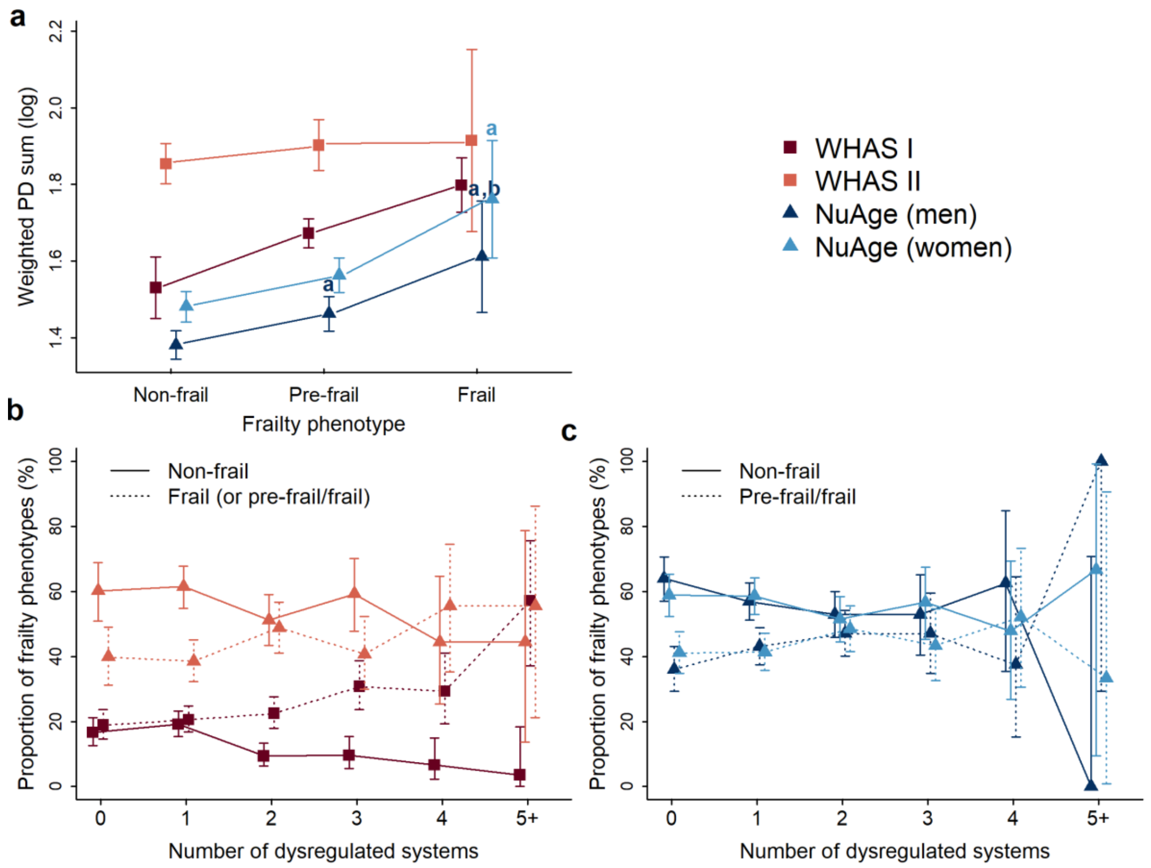


**Figure S6. Cumulative effect of the number of dysregulated systems on frailty phenotype, by data subset. (a)** Mean levels of a weighted sum of PD across six (WHAS I and WHAS II) and five (NuAge) systems by frailty phenotypes. Linear regressions were performed with frailty phenotype as a fixed effect, controlling for age with a cubic spline and for individual as a random effect. Lower letter "a" indicates significantly different from the non-frail phenotype, while lower letter "b" indicates significantly different from the pre-frail phenotype. **(b)** Prevalence of frail (dotted lines) and non-frail (solid lines) individuals according to the number of dysregulated systems in WHAS (red) and NuAge (blue) cohorts. Because the frail phenotype represents only ~3% of all observations in NuAge and WHAS II, we combined pre-frail and frail subjects for these cohorts. Different colors indicate different datasets (or subsets).

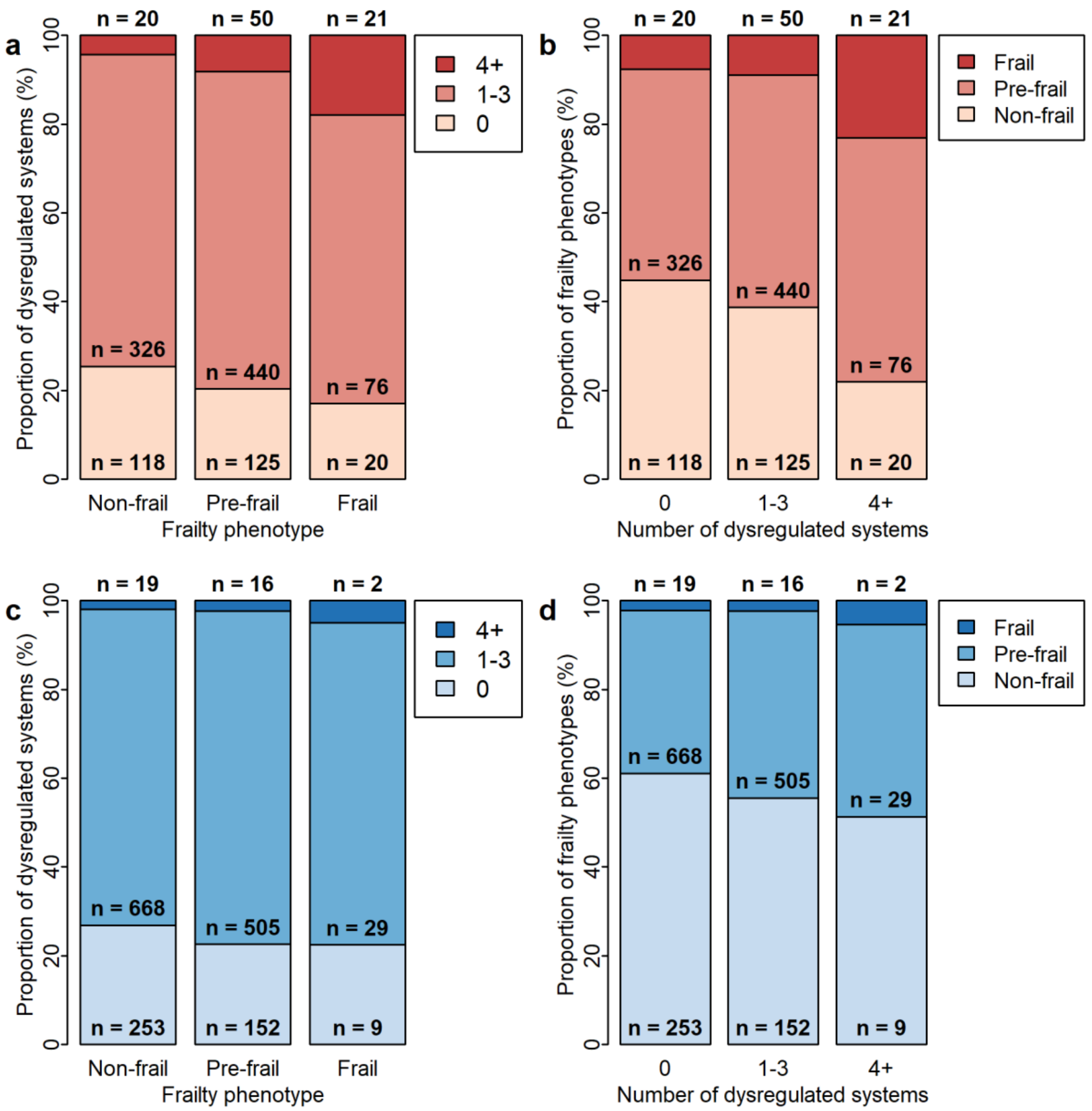


**Figure S7. Percentage of number of dysregulated systems by frailty phenotype, and vice versa.** The figure shows prevalences of dysregulated physiological systems according to frailty phenotypes in WHAS I + II (**a**) and NuAge (**c**), and prevalences of frailty phenotypes according to the number of dysregulated systems in WHAS I + II (**b**) and NuAge (**d**). The number of observations for each frailty phenotype and number of dysregulated systems combination is indicated.

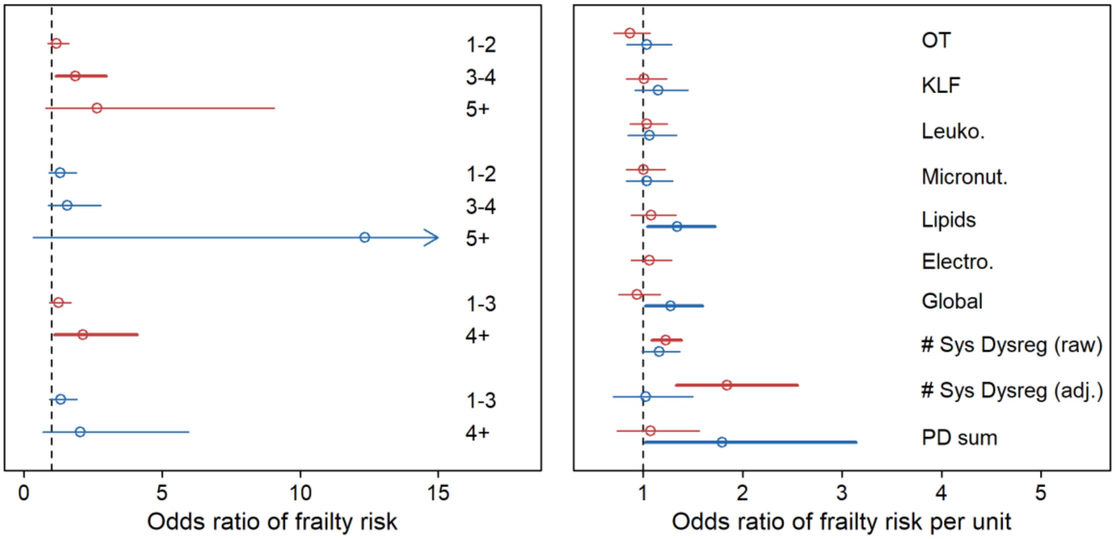


**Figure S8. Association between physiological dysregulation and frailty risk in full regression models, grouping pre-frail with frail** Estimations (points) together with 95% CIs (segments) are plotted for relationships between dysregulation levels, or the number of dysregulated systems ("# Sys Dysreg"), and frailty risk in WHAS (n = 1194, red) and NuAge (n = 1653, blue). **(a)** Frailty risk was assessed with a logistic regression model comparing non-frail to pre-frail/frail, as a function of the number of dysregulated systems categorized either as 1-2, 3-4, and 5+ (upper part) or 1-3 and 4+ (lower part), with no dysregulated system as the reference group. **(b)** Frailty risk was assessed with a logistic regression model comparing pre-frail/frail to non-frail, as a function of the number of dysregulated systems. All models excepted "# Sys Dysreg (raw)" controlled for the presence or absence of dysregulation in individual systems (dummy variables). Abbreviations: # Sys Dysreg, Number of dysregulated systems; Electro., Electrolytes; KLF, Kidney/liver function; Leuko., Leukopoiesis; Micronut., Micronutrients; OT, Oxygen transport.

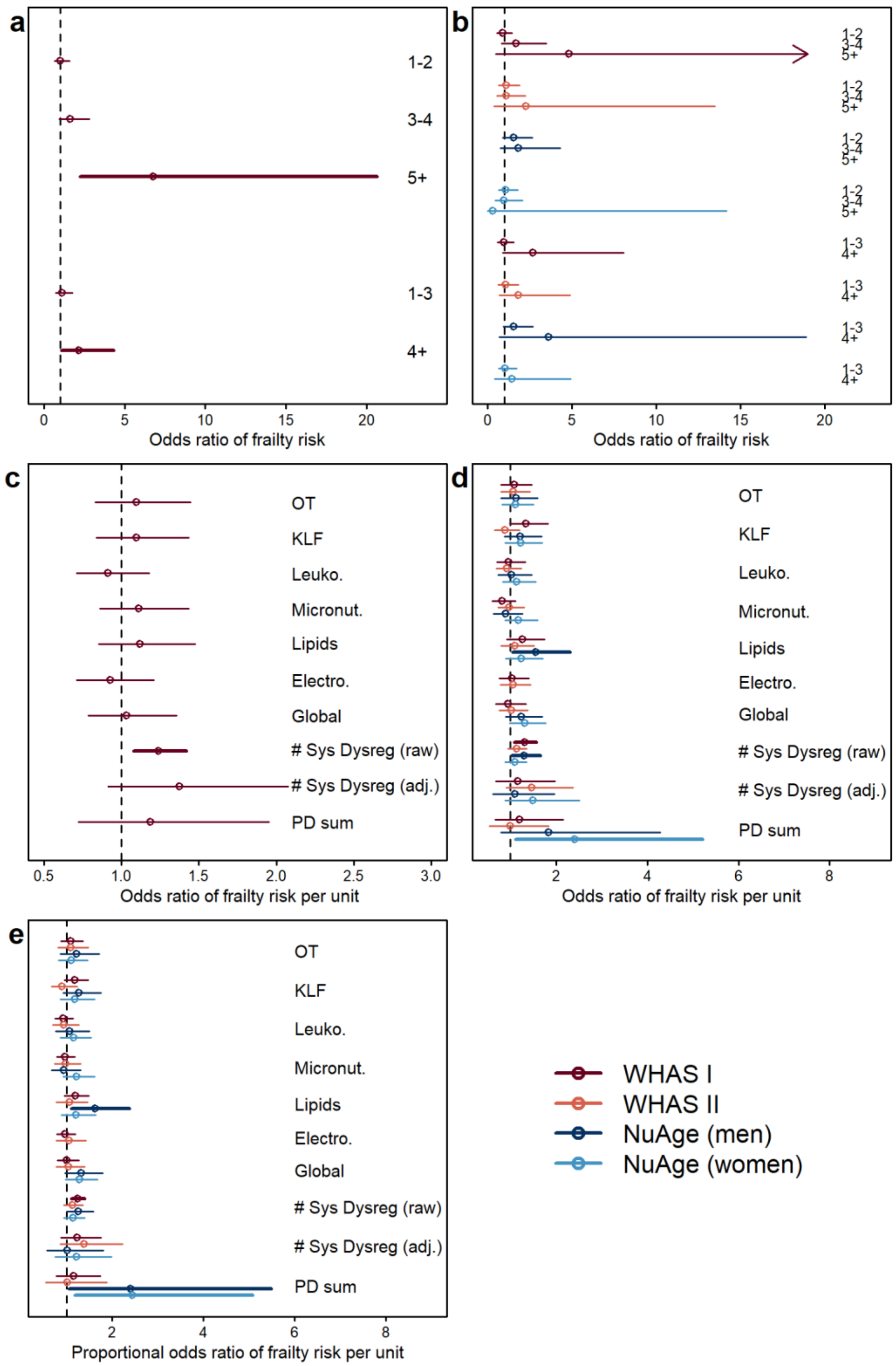


**Figure S9. Association between physiological dysregulation and frailty risk in full regression models, by data subset.** Estimations (points) together with 95% CIs (segments) are plotted for relationships between dysregulation levels, or the number of dysregulated systems ("# Sys Dysreg"), and frailty risk in WHAS I (n = 1277), WHAS II (n = 637), and in NuAge by sex (n = 795 and 858, respectively for men and women). **(a-b)**

Frailty risk was assessed with a logistic regression model comparing non-frail/pre-frail to frail (**a**) and non-frail to pre-frail/frail (**b**), as a function of the number of dysregulated systems categorized either as 1-2, 3-4, and 5+ (upper part) or 1-3 and 4+ (lower part), with no dysregulated system as the reference group. **(c-d)** Frailty risk was assessed with a logistic regression model comparing (**c**) non-frail/pre-frail to frail, and (**d**) pre-frail/frail to non-frail, as a function of the number of dysregulated systems. (**e**) Frailty risk was also assessed with a proportional odds model on non-frail to pre-frail to frail. All models excepted "# Sys Dysreg (raw)" controlled for the presence or absence of dysregulation in individual systems (dummy variables). Results are only shown for WHAS I when combining pre-frail to non-frail, because WHAS II and NuAge have very few frail individuals (~3% of individuals, see Table 1). Abbreviations: # Sys Dysreg, Number of dysregulated systems; Electro., Electrolytes; KLF, Kidney/liver function; Leuko., Leukopoiesis; Micronut., Micronutrients; OT, Oxygen transport.
